# Bipartite functional fractionation within the neural system for social cognition supports the psychological continuity of self *vs.* other

**DOI:** 10.1101/2021.04.04.438408

**Authors:** Rocco Chiou, Christopher R. Cox, Matthew A. Lambon Ralph

## Abstract

Research of social neuroscience establishes that regions in the brain’s default network (DN) and semantic network (SN) are engaged by socio-cognitive tasks. Research of the human connectome shows that DN and SN regions are both situated at the high-order end of cortical gradient but differ in their positions on this gradient. In the present study, we integrated these two bodies of research, used the psychological continuity of self *vs.* other as a ‘test-case’, and used fMRI to investigate whether these networks would encode social concepts differently. We found a robust dissociation between the DN and SN – while both networks contained sufficient information for decoding broad-stroke distinction of social categories, the DN carried more generalisable information for cross-classifying across social distance and emotive valence than did the SN. We also found that the overarching distinction of self *vs.* other was a principal divider of the representational space while social distance was an auxiliary factor (subdivision, nested within the principal dimension), and this representational landscape was more manifest in the DN than in the SN. Taken together, our findings demonstrate how insights from connectome research can benefit social neuroscience, and have implications for clarifying the two networks’ differential contributions to social cognition.

## Introduction

Decades of research suggests that humans deduce other individuals’ mental states by simulating how oneself would think and feel in similar situations (e.g., Steinbeis 2016). This ability is rooted in the awareness that self and others are distinct yet relatable social beings. The rudimentary sense of ‘self *vs.* other’ emerges during infancy, while self-concept (a sophisticated understanding of ourselves in relation to others under social contexts) evolves over lifespan. Neuroimaging evidence (for review, see Yeshurun *et al*. 2021) has established that the brain’s default-mode network (DN) is robustly engaged by various self-referential processes. This leads to the view that the DN is the key neural substrate that underpins ‘core self’/‘ego’ (Carhart-Harris and Friston 2010). Rather than a homogenous structure, converging results from seed-based connectivity, data-driven parcellation, and task-induced activation have further indicated that the DN comprises functionally distinct and anatomically separable subnetworks (e.g., Braga and Buckner 2017; Braga *et al*. 2019; Chiou *et al*. 2020). The division within the DN into subnetworks begs an important question – whether and how each subnetwork differentially contributes to the representation of self. In the present study, we investigated this issue by testing howthepsychological continuity from ‘self-ness’ to ‘otherness’ is encoded in two major subnetworks of the DN using multiple-voxel pattern analysis (MVPA). Our findings unravelled a systematic, graded pattern wherein the distributed neural coding for different social beings varied between brain regions and psychological constructs. Below we first discuss the neural basis of self-referential processes and the limitation of univariate approach. Next, we discuss evidence for a bipartite fractionation within the DN. We specifically focus on how such a fractionation constrains our scope of scrutiny for functional neural substrates, and how we can exploit MVPA to investigate the neural coding for ‘self *vs.* other’ within this bipartite structure.

Self-concept is a collection of beliefs about oneself, embodying the contents of contemplation about ‘*Who am I?*’ (Oyserman *et al*. 2012). It is inextricably intertwined with many critical issues in cognitive and social psychology, including perception of the aesthetics and attractiveness of one’s own body (self-image), the capability for reflecting on one’s dispositions (self-knowledge), one’s recollection of life events (autobiographical memory), etc. Previous research has shown that self-concept depends, in a fundamental way, upon one’s capacity to represent self as a psychologically coherent entity persisting through time, whose *past self* is construed as an entity closely related to yet partially separable from the *present self* (Klein *et al*. 2002). In addition to representing *present self* as a continuation of *past self*, research has shown that self-concept is impacted by the presence of other people in a social environment. Especially, during childhood and adolescence, self-concept is formulated through interaction with significant others, which spawned numerous investigations of familial and peer influences on personality and self-esteem (e.g., Deković and Meeus 1997; Verschueren *et al*. 2012). The significance of self-related processing is not only seen in the field of social psychology but also in cognitive neuroscience. Researchers have long understood that self-related processing is one of the major neurocognitive dimensions that characterise the functionality of the cortical midline structures, particularly the ventromedial prefrontal cortex (vmPFC) (for review, see Lieberman *et al*. 2019). For example, compared to information regarding another individual, processing information with reference to self amplifies vmPFC activity. This has been robustly observed when one evaluates descriptions with respect to self (Kelley *et al*. 2002), recalls personal experiences (Spreng and Grady 2010), or even recognises one’s own name/address that is ostensibly unrelated to the task at hand (Moran *et al*. 2006). In addition, patients with vmPFC lesion fail to show self-referential memory advantage (superior mnemonic performance for information related to self that healthy people reliably show), despite those patients having otherwise intact memory for non-self information (Philippi *et al*. 2012). Research also indicates that the vmPFC is sensitive to social distance with regard to self, with friends eliciting greater vmPFC activity than strangers, presumably due to one’s construal of a ‘friend’ being inextricably linked to oneself (Krienen *et al*. 2010). In a similar vein, vmPFC activity was found to reflect whether someone’s political opinion is in agreement with oneself, with encountering someone holding concordant views eliciting greater vmPFC activity than someone holding opposite views (Mitchell *et al*. 2006). Together, multiple threads of evidence consistently suggest that the vmPFC is a key neural structure underlying self-related processing, and is involved in processing other individuals when they are deemed similar/related to self.

The univariate approach is useful for identifying the loci of maximally-responding clusters/regions for a certain cognitive process, giving a bird’s eye view about where the ‘hotspot’ is. In the case of self-related processing, the ‘subtraction’ paradigm is often used – the univariate response of each voxel (or averaged response within a region of interest) elicited by self-related processing is compared with the response driven by evaluating, imagining, or reminiscing about someone else. While this approach is useful in identifying conspicuous targets in the brain (i.e., contiguous voxels that form a cluster surpassing thresholds), it has limited use in unveiling nuances that are jointly encoded by distributed patterns of neural responses across voxels. With MVPA, however, researchers are able to reveal how distributed neural patterns represent the subtle distinction between self- and other-referential processes. For example, the patterns of vmPFC activity have been used to decode if one was entertaining self- or other-referential thoughts (e.g., Chavez *et al*. 2016; Yankouskaya *et al*. 2017). In addition to deciphering self *vs.* other using vmPFC patterns, MVPA has been applied to decode various facets of social cognition using neural patterns elsewhere in the brain, such as decoding the identities of fictitious characters (Hassabis *et al*. 2013), the identities of personally familiar people (Thornton and Mitchell 2017), and whether a behaviour shows goodwill or malice (Koster-Hale *et al*. 2013). Together, these studies demonstrate how MVPA can be used to elucidate our understanding about the neural basis of social cognition, capturing the granularity in collective, distributed codes that often goes unnoticed by the univariate analysis (for review of recent progress, see Wagner *et al*. 2019).

While the vmPFC is heavily studied, it is not the only region involved in self-related processing. Alongside vmPFC activity, a group of widely distributed brain regions, which collectively form the brain’s specialised system for social cognition, have been reliably found to be involved in various self-related/socio-cognitive processes (for review, see Doré *et al*. 2014). For instance, vmPFC activity usually elevates in tandem with activity of other midline structures – the posterior cingulate cortex (PCC), retrosplenial cortex (RSC), and dorsomedial prefrontal cortex (dmPFC) – when a task requires assessing the personality of people, retrieving autobiographical episodes, or envisaging prospective and counterfactual scenarios that involve human activity. Apart from the brain’s medial aspect, self-referential/social-cognitive tasks also recruit a set of regions on the brain’s lateral surface (Olson *et al*. 2013; Doré *et al*. 2014; Binney *et al*. 2016; Chiou *et al*. 2020), including the bilateral anterior temporal lobes (ATL), the left inferior frontal gyrus (IFG), and the inferior parietal lobule (IPL). Resting-state research has shown that these areas, conjointly as the system for social cognition, tend to stay intrinsically connected during task-free resting moments, and overlap substantially with the cortical realm of the DN. Regions implicated in social cognition exhibit a similar functional profile that has been used to characterise the tendencies of DN regions (Raichle *et al*. 2001; Yeshurun *et al*. 2021): Akin to the DN, social regions are more active when a situation necessitates integrating internally-constructed representations (e.g., memories or schemas) with external signals (e.g., Murphy, Poerio*, et al.* 2019). Also resembling the DN, activation of the social system abates when a situation requires externally-oriented sensorimotor processes that minimally involve any internal representations (e.g., Chiou *et al*. 2020). Importantly, converging evidence from multiple investigations have shown that the extensive DN can be fractionated into (at least) two subsystems (Humphreys *et al*. 2015; Braga and Buckner 2017; Braga *et al*. 2019; Jackson *et al*. 2019; Chiou *et al*. 2020). While the nomenclature used to name the subsystems of DN varies between studies, a bipartite structure has been reliably observed in the profiles of different regions’ connectivity alliance and task-driven reaction, partitioning the DN into two modules. One subsystem comprises the vmPFC, the RSC, the posterior part of IPL, and the hippocampal formation. Some nodes of this subsystem are suggested to be the ‘hub’ nodes of the DN (Andrews-Hanna *et al*. 2010), and they are altogether dubbed ‘Network-A’ by more recent studies (e.g., Braga and Buckner 2017). These regions show a strong propensity to be *suppressed* by contexts that require externally-focused sensory-motoric processes (and *activated* by introspective processes), and are the quintessential ‘task-negative’ areas by traditional views of the DN (Raichle *et al*. 2001). The other subsystem consists of the bilateral ATL, the left IFG, and temporoparietal junction. In the field of semantic cognition, these regions are dubbed the ‘semantic network’ (SN; owing to their robust activity during semantic processes), while in the field of connectome research, they are dubbed ‘Network-B’. While many areas of the SN are incorporated under the umbrella of DN regions (Yeo *et al*. 2011), recent evidence has indicated that the SN is a functionally separable entity from DN. Specifically, DN and SN regions show a similar inclination to deactivate in outwardly-oriented sensorimotor tasks (although the extent of ‘aversion’ to sensorimotor processing is significantly more moderate in SN regions, see Chiou *et al*. 2020). However, the two subnetworks diverge on reaction to semantic processing – whereas SN activity intensifies for verbal and non-verbal semantic processing, DN activity attenuates (Humphreys *et al*. 2015; Chiou *et al*. 2018; Jackson *et al*. 2019). The dissociation between the two subnetworks was robust, found both in resting-state and task-based fMRI studies. Together, multiple lines of inquiries have consistently demonstrated that the DN and SN are functionally distinct entities (whilst both are heavily engaged by social cognition). It is noteworthy that there is less consensus on the taxonomy of the dmPFC. The dmPFC shares a similar functional profile to other DN regions during a variety of social-cognitive tasks (Hiser and Koenigs 2018), and is reliably linked with other core nodes of the DN, particularly the vmPFC and PCC/RSC (e.g., Bzdok *et al*. 2015; Eickhoff *et al*. 2016; Jackson *et al*. 2020). However, the dmPFC is also affiliated with the SN – relative to the vmPFC as a seed, the dmPFC has tighter functional coupling with many SN regions in resting-state (Bzdok *et al*. 2013), and is more reactive to tasks that lay emphasis on extracting semantic meaning from lexical stimuli than other DN regions (Chiou *et al*. 2020). Pertinent to our main question, while the DN and SN dissociate on their univariate reaction to different contexts, it remains unclear whether the two subnetworks carry distinct fine-grained multiple-voxel patterns that convey different information about self *vs.* other.

In the present study, we investigated the distributed neural coding that underpins the continuity of ‘self *vs*. other’ representations. Self-concept is known to evolve along one’s lifetime. However, although one’s beliefs about *present self* might be radically different from his/her opinions about *past self*, the human mind maintains the stability/continuity of self-identity, treating *present self* and *past self* as dissociable yet associated entities that represent the same person (Northoff 2017). Psychological continuity is also applicable to the construal of self *vs.* others, with a close other (or personally familiar person) perceived as more affiliated with (similar to) self than a distant other (personally unrelated person). It remains unknown how such continuity is encoded in the brain. We instigated this issue by gradationally manipulating social distance. We established four points of reference: the participant’s sense of *present self*, the participant’s sense of *past self* ten years ago, a personally familiar other – the participant’s mother, and a personally unfamiliar but well-known other – Queen Elizabeth II. Behavioural rating confirmed that participants rated the four persons along a continuum, with *present self* being most distinct from *the Queen* while *past self* and *mother* situate serially somewhere in between. While undergoing fMRI, participants read descriptions (either positive or negative personality traits) in contexts of each of the four individuals and judged whether the description aptly characterised the person. We employed two types of MVPA, machine-learning classification (Pereira *et al*. 2009) and representational similarity analysis (Nili *et al*. 2014), to investigate how the two subnetworks encoded the psychological continuity of self *vs.* other and the possible divergence between DN and SN.

Replicated across multiple analyses, we found (*i*) the broad-stroke division of self *vs.* other was a principal axis of the representational space while social distance was an auxiliary axis, nested within the principal dimension; (*ii*) the DN dissociated from the SN, with the former carrying more information about personal identities. In the section of Discussion, we expound on how recent progress in understanding the topography of cortical mantle (particularly studies that demonstrated that the DN is the apex of cortical hierarchy; Margulies *et al*. 2016) could be leveraged to explain the dyadic split between the DN and SN and their differential contributions to cognition.

## Materials and Methods

### Participants

Twenty-four volunteers gave informed consent before the fMRI experiment. The sample consisted of a 10/14 female-to-male ratio, with average age = 33 years-old and SD = 11. All volunteers are right-handed and speak English as their mother tongue. All of them completed the magnetic resonance imaging safety screening questionnaire before the experiment and reported not having any neurological or psychiatric condition. This study was reviewed and approved by the local research ethics committee.

### Experimental design

Participants completed two experiments while undergoing fMRI in a single session. In the main experiment, participants read short phrases describing various traits of personality or temperament, either positive or negative, and made a binary button response to answer whether they reckoned the phrase rightly described the characteristics of the specific person that was under consideration. In the localiser experiment, participants read short narratives describing either events that involved human activity or events that involved alterations of the physical environment, and made a binary button response to answer a comprehension question related to the narrative they just read. A session began with the acquisition of each participant’s anatomical scan, followed by the main experiment that consisted of eight functional runs of scanning, and the localiser experiment that contained only a single run.

The main experiment had a 4 × 2 factorial design (four individuals: *Present Self*, *Past Self*, *Mother*, and *the Queen Elizabeth II*; two sides of emotive valence: positive *vs.* negative traits). We adopted and modified a well-established fMRI paradigm that has been widely used to assess the neural substrates of self- and other-referential processing (e.g., Kelley *et al*. 2002; Heatherton *et al*. 2006; Meyer and Lieberman 2018). The task was to read a short description in each trial and to evaluate whether or not it appropriately depicted a particular individual. When performing the task, participants were presented with a fixation dot (0.5 sec) in each trial, followed by words (3.3 sec). They were required to make a response within the 3.3-sec time-limit. The target of assessment (*Present Self*, *Past Self*, *Mother*, and *Queen*) was shown above the fixation dot, and a short phrase describing a certain personality or temperament trait was shown blow (e.g., ‘*Sincere to friends*’ or ‘*Anxious about uncertainty*’). When the target was *Past Self*, participants reflected on themselves specifically ten years ago with respect to the phrase. Stimuli were presented using a block design, controlled with E-Prime (Psychology Software Tools). Each run consisted of 16 blocks of trials, with each of the eight conditions having two blocks. Across the runs and participants, the order in which the eight task-conditions were presented was counterbalanced so that each task-condition was equally likely to appear in each of the 128 possible slots of the eight runs (i.e., each condition was equally probable to preceded or succeed any other condition), with stimuli randomly drawn from a designated stimuli-set for a given run and shuffled across blocks. The stimuli-sets were also counterbalanced across participants; thus, each set was equally likely to be presented in each run. This fully counterbalanced design is vital for the subsequent leave-one-run-out decoding analysis, ensuring that every task-condition and stimuli-set appeared in each fold of the cross-validation. Each block was 19-sec long, containing five trials and no inter-trial interval. Each run of scanning was 380-sec long, containing 16 task-blocks, 15 inter-block intervals (blank screen, 5-sec each), and a 1-sec blank at the end. All text stimuli were white in colour, Arial typeface, 28-point in font size displayed on a black background. Participants reacted to the questions by pressing one of the two designated buttons on a MR-compatible response-pad with their right index or middle finger. All visual stimuli were displayed using high-resolution LCD goggles (NordicNeuroLab) mounted on top of the head coil.

A total of 160 short phrases were used for evaluating personality traits. Each phrase contained three or four words. A half of the phrases were designed to convey positive meaning, whereas the remaining half conveyed negative meaning (the complete set of stimuli are reported in Supplementary Material). To ascertain that the phrases express the emotive meaning as intended, we asked six volunteers (none participated in the later fMRI study) to rate the emotive valence of the phrases using a 5-point scale (1 being most negative, 5 being most positive). Results of rating support the adequacy of our stimuli: By-subject analysis showed that phrases designed to convey negative meaning (average±SEM: 1.7±0.1) were rated significantly lower than those designed to convey positive meaning (average±SEM: 4.4±0.1; *t*_(5)_ = 14.5, *p* < 0.0001). This difference was also seen in by-item analysis for every volunteer (all *p*s < 10 ^-10^). The length of text stimuli was also equated: No difference was found between the letter counts between negative (average±SD: 22±3.4) and positive phrases (average±SD: 21±3.8, *p* > 0.21, *n.s.*). Crucially, each of the 160 phrases was equiprobable to be assessed with reference to any of the four target individuals. Namely, every phrase appeared four times in the experiment but referred to a different individual each time it was presented. This circumvented the potential biasing effect of specific stimuli by equating their frequency in every condition, and ensured that none of the stimuli was repeated – each combination of description and person was encountered only once during the experiment and appeared ‘novel’ from a participant’s perspective. This set-up also ensured that identical stimuli (descriptions of personality) were used in each condition of the four target individuals, with the only difference being the cue word that reminded the current target.

The localiser experiment was based on a well-established paradigm that has been repeatedly used to assess the brain regions associated with the processes of mentalising/theory of mind (e.g., Saxe and Kanwisher 2003; Saxe and Wexler 2005; Dodell-Feder *et al*. 2011). There were two conditions in this localiser run: In the Social condition, participants read a narrative describing human interactions; afterwards, they were presented with a statement about the beliefs or feelings that a person in the narrative might have, and were asked to verify whether the statement is true or false. Answering such questions entailed changing perspectives and making inferences about someone’s mental states. In the Non-social condition, participants read a narrative describing the physical state of the world (e.g., “*a large oak tree stood in front of the city hall from the time the building was built…*”) and answered a comprehension question related to the non-social narrative. Participants had 10 sec to read the narrative and were probed with an ensuing question that was shown for four sec. The localiser run was 520-sec long, containing 20 task-blocks (10 Social blocks, 10 Non-social blocks) and 20 blank intervals (each 12-sec long) that followed each task-block. The two conditions were presented in an alternating order, and counterbalanced across participants (i.e., a half of them started with the Social condition; the other half started with the Non-social). We used the same text stimuli previously used by Dodell-Feder *et al*. (2011). We adopted this localiser task due to the fact that, in the original study, this paradigm had proved effective in reliably activating a set of widely distributed brain regions known to be sensitive to mentalisation in specific (and social cognition in general), which incorporates all of our regions of interest (ROI) in the default-mode network (DN), as well as in the semantic network (SN; see the details of how we defined the voxels in the section of Regions of Interest).

### MRI acquisition

Some of the regions of our primary interest are situated in the rostro-ventral aspects of the brain (e.g., the ATL and the vmPFC), which are known to be particularly susceptible to signal-dropout issues (Visser *et al*. 2010). To combat signal-dropout in these areas, we adopted a dual-echo EPI sequence, which has been demonstrated to effectively improve signal-to-nose ratio in dropout-prone regions, compared to other conventional imaging protocols (for precedents using this dual-echo acquisition protocol, see Halai *et al*. 2014; Jackson *et al*. 2015; Chiou and Lambon Ralph 2019). Scans were acquired using a 3T Phillips Achieva scanner equipped with a 32-channel coil and a SENSE factor of 2.5. Using this protocol, each scan consisted of two images acquired simultaneously with two echo-times: a short echo optimised to obtain maximum signal from the ventral parts and a long echo optimised for whole-brain coverage. The sequence included 31 slices covering the whole brain with repetition time (TR) = 2.8 sec, short/long echo times (TE) = 12/35 ms, flip angle = 85°, field of view (FOV) = 240 × 240 mm, resolution matrix = 80 × 80, slice thickness = 4 mm, and voxel dimension = 3 × 3 mm on the x- and y-axis. To reduce ghosting artefacts in the temporal lobes, all functional scans were acquired using a tilted angle, upward 45° off the AC-PC line. For the main experiment, the EPIs were collected over eight runs; each run was 380-sec long during which 136 dynamic volumes were acquired (alongside 2 dummy scans, discarded). For the localiser experiment, the EPIs were collected from a single run (520 secs); 186 dynamic volumes (and 2 dummies) were acquired. To tackle field-inhomogeneity, a B_0_ field-map was acquired using identical parameters to the EPIs except for the following: TR = 599 ms, short/long TEs = 5.19/6.65ms. Total B_0_ scan time was 1.6 minutes. A high-resolution T1-weighted structural scan was acquired for spatial normalisation (260 slices covering the whole brain with TR = 8.4 ms, TE = 3.9 ms, flip angle = 8°, FOV = 240 × 191 mm, resolution matrix = 256 × 163, and voxel size = 0.9 × 1.7 × 0.9 mm).

### Preprocessing and GLM

An established procedure was used to combine the two volumes from the dual-echo dataset. Using SPM8 (Wellcome Department of Imaging Neuroscience), we integrated the standard pre-processing procedure (realignment, slice-time correction, co-registration, and the linear integration of the long- and short-echo images) with B_0_ field-map correction to prevent distortion due to inhomogeneity. The linear averaging approach has been well-established in previous studies (e.g., Poser *et al*. 2006; Halai *et al*. 2014; Chiou *et al*. 2018). The combined images were realigned using rigid body transformation (correction for motion-induced artefacts) and un-warped using B_0_ field-map (correction for field-inhomogeneity). The integrated EPIs were then co-registered with each participant’s T_1_ anatomical image. For the 1^st^- level individual analysis, the *β*-weight of each experimental regressor was estimated by convolving each task-block with a canonical haemodynamic response function. Six motion parameters were added into the model as nuisance covariates in the general linear model. Behavioural reaction times were also modelled as parametric modulators to account for the influence of fluctuating reaction times within a condition. For the main experiment, each of the eight experimental conditions were modelled explicitly as a separate regressor, while resting baseline was modelled implicitly. For the localiser experiment, the entire 14-sec duration of ‘story’ and ‘question’ intervals was convolved with a canonical hemodynamic response function, as per previous studies using the same localiser paradigm (e.g., Dodell-Feder *et al*. 2011; Skerry and Saxe 2014, 2015). Low-frequency drifts were removed using a high-pass filter of 128 sec. Beta-estimates associated with each voxel and each regressor were subsequently submitted to multivoxel pattern analysis.

Decoding analysis was performed on each participant’s brain in native space, without normalisation and smoothing. Normalisation (MNI) and smoothing (FWHM = 8mm) were only done on the individual whole-brain searchlight outcomes (accuracy maps), prior to the 2^nd^-level random-effect analysis. The decoding accuracy maps of whole-brain searchlight analysis were normalised into the MNI standard space using the DARTEL Toolbox of SPM (Ashburner 2007), which has been shown to produce highly accurate inter-subject alignment (Klein *et al*. 2009). Specifically, the T_1_-weighted image of each subject was partitioned into grey-matter, white-matter, and CSF tissues using SPM8’s ‘Segmentation’ function; afterwards, the DARTEL toolbox was used to create an average template combining all participants of the group. The grey-matter component of this template was registered into the SPM’s grey-matter probability map (in MNI) using affine transformation. In the process of creating the group’s template using individual T_1_, for each individual DARTEL estimated ‘flow fields’ that contained the parameters for contorting native T_1_-weighted images to the group template. SPM8 deformation utility was then applied to combine group-to-MNI affine parameters with each participant’s ‘flow fields’ to enable tailored warping into the standard MNI space. At the end of the procedure, the parameters of transformation were applied to the whole-brain map of searchlight-decoding data, and resampled voxel size using 3 × 3 × 3 mm. Smoothing on the normalised accuracy maps was then applied using an 8-mm Gaussian FWHM kernel, consistent with prior studies (e.g., Halai *et al*. 2014; Jackson *et al*. 2015).

### Regions of interest

As discussed earlier, we adopted a localiser paradigm that had previously proved effective in detecting various target regions in the DN and SN. In the original study that reported this fMRI paradigm, Dodell-Feder *et al*. (2011) tested a sizable sample of 62 participants and contrasted the Social condition against the Non-social one. Using this contrast, they detected robust activation in five areas of the DN – the dmPFC, vmPFC, PCC, and left/right IPL, as well as in three areas of the SN – the left/right ATL and IFG. Because the Dodell-Feder *et al*. results provided a useful exemplar regarding the loci of neural activity, their group-level peak coordinates (in the MNI space) could be used to guide the localisation of ROIs in our participants’ native space. For each participant’s localiser data, we applied a *t*-test, voxel-wise thresholded at *p* < 0.001, to generate a whole-brain map of *t*-values to identify voxels that responded more intensely to the Social than Non-social condition. Using each participant’s reversed-normalisation parameters computed by SPM, we first identified eight ‘landmark’ points (5 DN regions, 3 SN regions) in each person’s native brain that corresponded to the coordinates from the Dodell-Feder *et al*. study. Next, we identified the local maxima nearest to each of the eight ‘landmark’ points and created a spherical ROI (radius = 10 mm) centred at the local maximal coordinate. If no activity was detected at *p* < 0.001, we repeated the procedure using *p* < 0.005 and *p* < 0.01. This procedure constrained the localisation of ROIs using the group-level results from Dodell-Feder *et al*. (2011) while allowing subject-specific variation in functional activation. For every participant we were able to identify eight ROIs – five regions of the DN (the dmPFC, vmPFC, PCC, and left/right IPL) and three regions of the SN (the left/right ATL, and IFG). Meticulous care was taken to ensure that all ROIs were spatially mutually exclusive – namely, there was no overlap in the voxels contained in each functionally-defined ROI (particularly for the dmPFC and vmPFC). It is worth emphasising that, with an independent localiser task to select the ROIs, our subsequent MVPA analyses were devoid of the statistically ‘double-dipping’ issue (Button 2019).

### Multivoxel pattern analysis

Prior to multivariate decoding, we fitted the main experiment’s data of each participant using a standard general linear model (GLM), implemented in SPM8, to compute the *β*-estimates of voxel-wise activation elicited by each experimental regressor. Beta-estimates were computed for each of the eight experimental conditions (*4* individuals: *Present Self*, *Past Self*, *Mother*, and *Queen* by *2* types of valence: Positive *vs.* Negative) and for the eight runs of scanning, yielding 64 *β*-weights. It is important to note that, for machine-learning classification, there is a trade-off between having many noisy examples (e.g., one *β-*value per trial) and having fewer but cleaner examples (e.g., one *β* for each task-condition per run – obtained by averaging examples of the same class to subdue their variability; for discussion on this issue, see Pereira *et al*. 2009; Haynes, 2015). Because our investigation concerned the very subtle representational differences between high-order concepts (e.g., one’s sense of self at present *vs.* in the past), the *β*- estimate of a single trial (or even the *β*-value computed for a single block) might introduce stimuli-specific noise and fail to reflect the most critical facet. Therefore, in order to de-noise, we prioritised having somewhat fewer but robust examples over many but flimsy examples by estimating a *β*-value per regressor per run. However, it is important to emphasise that, due to the factorial design that we employed, we still obtained sufficient examples for the training and testing – for instance, in the 4-way cross-classification wherein we decoded the four persons, trained using positive trials, and tested using negative ones (and *vice versa*), two separate sets of 32 examples were used for training and testing. Our dataset is sufficiently large for training and validation by the conventional practice of decoding research (Pereira *et al*. 2009). We performed MVPA with the Decoding Toolbox (TDT; Hebart *et al*. 2015), which employed a linear support-vector-machine (SVM) for classification with a ‘*C* = 1’ cost parameter (Chang and Lin 2011). Using standard approaches of cross-validation and cross-classification, we carried out six analyses to decode different aspects of ‘self-ness *vs.* otherness.’ For each classification, a leave-one-run-out eight-fold splitter was used whereby the algorithm learnt to distinguish between relevant categories using the data from seven of eight runs; its ability to correctly predict was tested using the unseen data from the remaining ‘held-out’ run. This procedure was iterated over all possible combinations of runs used for training and testing. By partitioning the datasets based on different runs of scanning, we ensured that there was no contamination of information leaking from the training-sets to testing-sets. The accuracy scores were then averaged across folds to produce a mean accuracy score for further statistical analysis. This was done separately for each participant, each ROI, and each of seven decoding analyses. The analyses included: (**1**) an aggregate sense of ‘self’ (incorporating *Present Self* and *Past Self*) *vs.* an aggregate sense of ‘others’ (incorporating *Mother* and *Queen*); (**2**) finer-grained differentiation within the self-concepts – *Present Self vs. Past Self*; (**3**) finer-grained differentiation within the concepts about others – *Mother vs. Queen*; (**4**) four-way differentiation amongst the four individuals – *Present Self vs. Past Self vs. Mother vs. Queen*; (**5**) Cross-classification of Self *vs.* Other across near and far social distances – the classifier was trained to tell apart self *vs.* other based on samples with closer social distance (i.e., *Present Self vs. Mother*) and tested using samples with farther distance (i.e., *Past Self vs. Queen*); this was repeated with the reverse mapping (i.e., using the ‘distant’ pair for the algorithm to learn and the ‘close’ pair to test its generalisability); (**6**) Cross-classification of the abstract sense of social distance across the self and other domain – the classifier was trained to distinguish *Present Self* from *Past Self* and tested using *Mother vs. Queen* (and *vice versa*). Successful cross-classification in this case indicated the acquisition of neural patterns that encoded the abstract information about near *vs.* far social relationship, applicable both to the self and other domain, (**7**) Cross-classification of four individuals across the two sides of emotive valence; the classifier was trained to perform four-way classification using the dataset wherein personality judgements were based on positive traits, and then was tested whether it could generalise to the unseen data based on negative traits. This cross-classifying was repeated with the inverse mapping (trained on the negative trials, tested on the positive ones). Bonferroni correction was applied to adjust multiple comparisons based on the number of ROIs in each analysis (α-level: 0.05/8 = 0.006).

As an exploratory analysis, we used the roaming whole-brain ‘searchlight’ method to test whether brain regions outside our selected ROIs also carried task-relevant information that allowed successful multivoxel-pattern decoding (Kriegeskorte *et al*. 2006). Each ‘searchlight’ was composed of a multivoxel pattern in a local neighbourhood (a sphere of 10-mm radius) that surrounded each voxel of the brain. For each sphere, a linear support-vector-machine classifier was trained to decode relevant information and tested using the same leave-one-run-out procedure as described above; the accuracy score was assigned to the centroid voxel. This process was repeated each time using a different voxel as the centre as the searchlight roamed across the brain. Prior to group-level analysis on the decoding result, each participant’s classification-accuracy map was normalised into the MNI space using the SPM’s DARTEL toolbox (the Pre-processing section) and smoothed using a Gaussian kernel of 8-mm FWHM. Voxel-wise decoding accuracy were tested against the chance-level (50% for binary classifications; 25% for four-way classifications) using a one-sample *t*-test. We constrained the rate of family-wise error (FWE) under *p* < 0.05 at the voxel-wise level to adjust for multiple comparisons.

As a complementary approach to machine-learning decoding, representational similarity analysis (RSA) was employed to investigate whether there was any systematic structure that underlay the neural representations as revealed by the pair-wise resemblance between different task-conditions (e.g., whether the multivoxel-patterns of *Present Self* are more similar to *Past Self* than to *Queen*). Based on the established methods (Kriegeskorte *et al*. 2008; Nili *et al*. 2014), we calculated the neural similarity between each pair of experimental conditions as the Pearson correlation of their vectorised patterns of voxel-wise activity. Note that we used similarity (Pearson’s *r* between patterns) as the metric, rather than their distance/dissimilarity (1 – Pearson’s *r*), to characterise the resemblance between contexts, for the sake of more intuitive and straightforward interpretations (the two approaches generated exactly the same conclusion as by mathematical definition they were two sides of the same coin; for discussion, see Dimsdale-Zucker and Ranganath 2018). Hierarchical clustering analysis was performed on the distance measures (1-*r* or 1*-*τ-a/*ρ*) to visualise the categorical clump/division between neural representations, embodied in the structure of a dendrogram tree. As per the standard approach of RSA research (e.g., Nili *et al*. 2014), we used the rank-based correlation indices – Kendall’s Tau (τ-a) and Spearman’s Rho (*ρ*) – to assess the 2^nd^-order relationship between two representational similarity matrices, and considered only the non-redundant/off-diagonal lower-triangular elements of each matrix to prevent inflation of correlation size. Given the non-parametric nature of τ-a and *ρ*, signed-rank test was used to assess whether the correlation between two representational similarity matrices was significantly greater than chance (one-tailed), as well as whether the correlation for a certain pair of matrices significantly differed from those of other matrices (two-tailed). Multiple comparisons were adjusted by constraining false-discovery rate (FDR) under 0.05. Finally, to evaluate the magnitude of correlation with reference to a hypothetically ‘true’ model’s optimal performance (given the amount of noise in the present data), the noise-ceiling was derived by (*i*) calculating the averaged correlation between the group’s matrix and every individual matrix (the upper-bound) and (*ii*) performing an iterative leave-one-subject-out procedure that correlated each individual’s matrix with all other participants’ averaged matrix (the lower-bound; *cf.* Nili *et al*., 2014).

## Results

### Machine-learning classification analysis

We sought to clarify (*i*) whether there is a spectrum-like change in neural coding in relation to social distance and (*ii*) whether or not the two subnetworks were equally capable of representing this ‘psychological continuity’. Clarifying these two research questions allowed us to decipher how the multivoxel patterns of DN and SN encoded the continuity and boundary of ‘self *vs.* other’ representations. In *Supplemental Results 1*, we report behavioural data and a confirmatory analysis of the motor cortex that refutes performance-related explanations. To localise regions of interest (ROI) unbiasedly in each participant’s native-space brain, we used a localiser task to independently identify five regions in the default-mode system and three regions in the semantic system. Using the peak-coordinates derived from the localiser data^1^, we defined eight spherical ROIs within the DN and SN – five ROIs are typically affiliated with the DN (the dmPFC, the vmPFC, the PCC, the left IPL, the right IPL), while three are associated with the SN (the left ATL, the right ATL, the left IFG). Support-vector machines (SVM) were trained on the neural pattern of each ROI. Using a supervised-learning cross-validation procedure, the algorithms were tested using ‘quarantined’ data to evaluate whether it was able to predict the mental content (i.e., a particular person one was reflecting on), and whether the chance of successful prediction varied systematically with social distance and network membership.

We began by verifying whether the regional multivoxel-pattern of each ROI allowed deciphering the broad-stroke information about the aggregate sense of self-ness (*Present Self* and *Past Self*) *vs*. the aggregate sense of otherness (*Mother* and *Queen*). Significantly above-chance decoding was achieved in every ROI (Figure 1A), suggesting that this coarse-grained, binary distinction between self and other is discernible using the patterns of local DN/SN activity. Motivated by this result, we tested whether it would be possible to decode nuance within the orbit of self-referential ideation (*Present Self vs. Past Self*) and other-referential ideation (*Mother vs. Queen*). As Figure 1A shows, statistically reliable decoding was achieved in nearly all ROIs (all, except for the left IFG) for the differentiation between *Present Self* and *Past Self*, and in every ROI for the differentiation between *Mother* and *Queen*. This indicates that the regional pattern of DN and SN activity enabled accurate classification not only *between* the domains of self *vs.* other, but also the finer-grained subgroups *within* the realm of self and other. Successful decoding of a socially proximal (*Present Self*) *vs.* a distant concept (*Past Self*) within the ‘self’ domain suggests that information about ‘social distance’ intersected with ‘self *vs.* other’. To untangle this intricacy, we performed four-way classification amongst the four individuals to understand how decoding fared when the algorithm had to consider ‘self *vs.* other’ and ‘social distance’ simultaneously. As shown in Figure 1A, while the chance-level dropped from 50% (binary) to 25% (four-way), reliable above-chance decoding was achieved in every ROI (also see *Supplemental Results 6* for the confusion matrices), indicating that the neural pattern that encoded each individual’s specific identity can be reliably differentiated from all other identities. Critically, a systematic trend was replicated across all of the four analyses: Although significantly above-chance decoding was obtained in every ROI, classification accuracy was reliably higher in the five ROIs that belong to the DN (with the PCC reliably reaching the highest accuracy), compared to the three ROIs that belong to the SN. This ‘imbalance’ between the two subnetworks implies that the DN, as a whole, carried greater amount of information about personal identities that could be extracted by the classifier to enable correct predictions. We also examined whether the univariate amplitude of each ROI could be used to differentiate individuals. As shown in Figure 1(B), results revealed that univariate contrasts did not reliably differ between conditions, suggesting that encoding personal identity relied on multivariate patterns rather than differential amplitudes at the univariate-level (also see *Supplemental Results 5* for the results of whole-brain research for univariate effects).

**Fig. 1.**
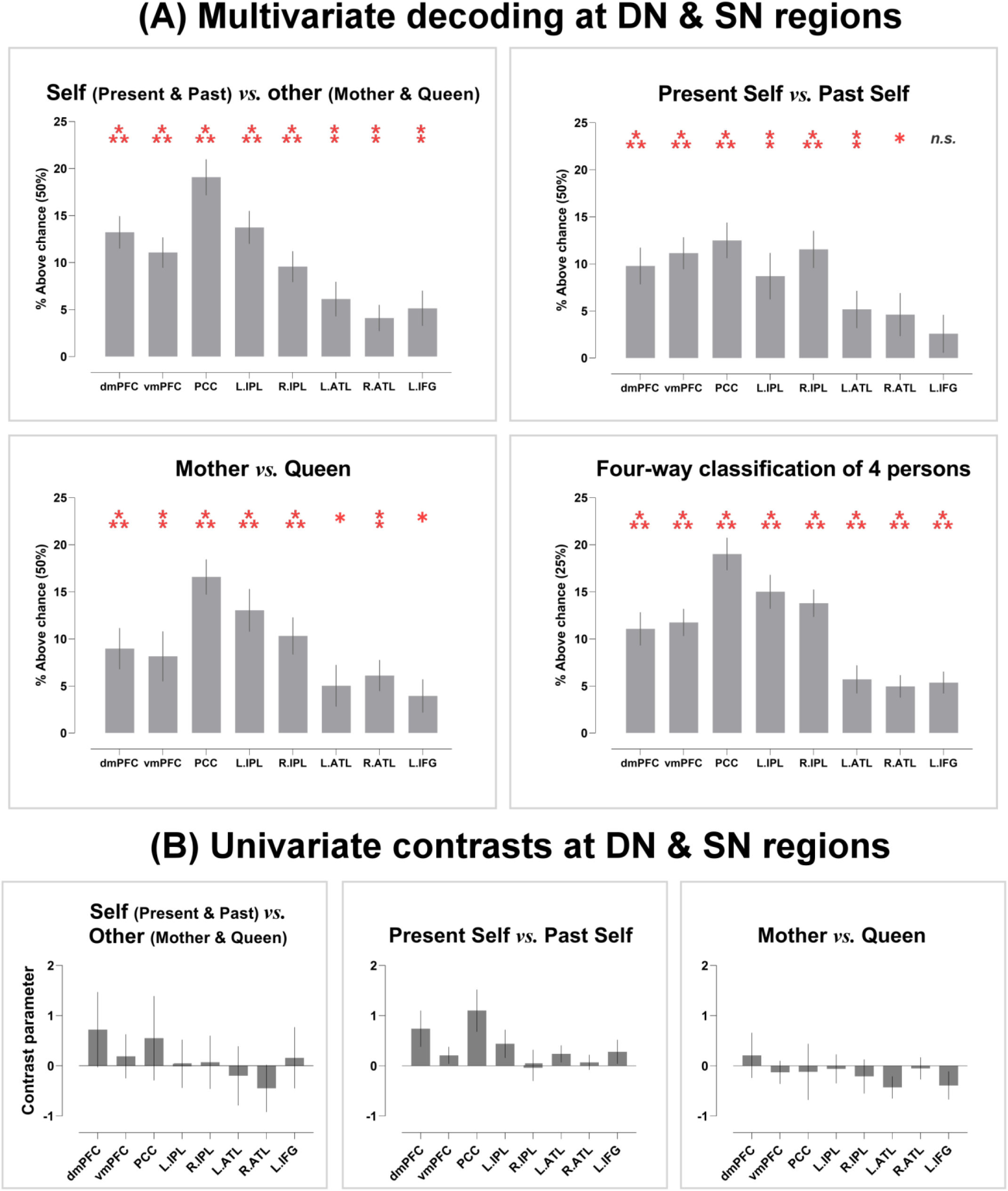
**(A)** Average accuracy of four classification analyses in the eight ROIs; asterisk indicate significantly above-chance decoding (Bonferroni-corrected for multiple comparisons). **(B)** Univariate contrast parameters in the same eight ROIs. **\*** *p* < 0.01; **\*\*** *p* < 0.005; **\*\*\*** *p* < 0.001.

Although identical stimuli were used in each condition (i.e., the same personality descriptions were assessed with reference with different individuals), participants saw different cue words that reminded them of the current target person. Thus, an alternative explanation is that our decoding was driven by word length – e.g., one word (Mother) *vs.* two words (Present Self). This alternative is unlikely given the fact that robust decoding was still achieved when the lengths of cue words were matched between conditions – see Figure 2(A) for the whole-brain searchlight interrogation of *Present Self vs. Past Self* and 2(B) for *Mother vs. Queen*. To further rule out the potential effects of low-level visual features (i.e., vertical/horizontal lines and curves that constituted the cue words), we trained the classifier to tell apart *Present Self vs. Past Self* and tested whether it could distinguish *Mother vs. Queen* (and *vice versa*). This cross-classification was a stringent test to assay whether the algorithm truly deciphered high-level information (in this case, social distance that was generalisable between self and other), independent of sensory factors (there was minimal visual resemblance between the pair of *‘Present Self vs. Past Self’* and *‘Mother vs. Queen’*). Results showed that cross-classification was detected in the default network – models trained to discriminate *Mother* from *Queen* could also reliably discern *Present Self* from *Past Self* (and *vice versa*). This suggests that the multivariate classifiers were, at least in part, overlooking sensory differences of letters and discovering neural patterns that enciphered social distance. As illustrated in Figure 2(C), successful cross-classification was achieved using the patterns of three DN regions in the midline structures (with the vmPFC being the most informative area), while decoding was at chance in two control regions (the primary visual and motor cortices). This rigorous verification suggests that DN regions carried information about personal identity and social proximity, and abstracted such information away from the sheer appearance of visual stimuli.

**Fig. 2.**
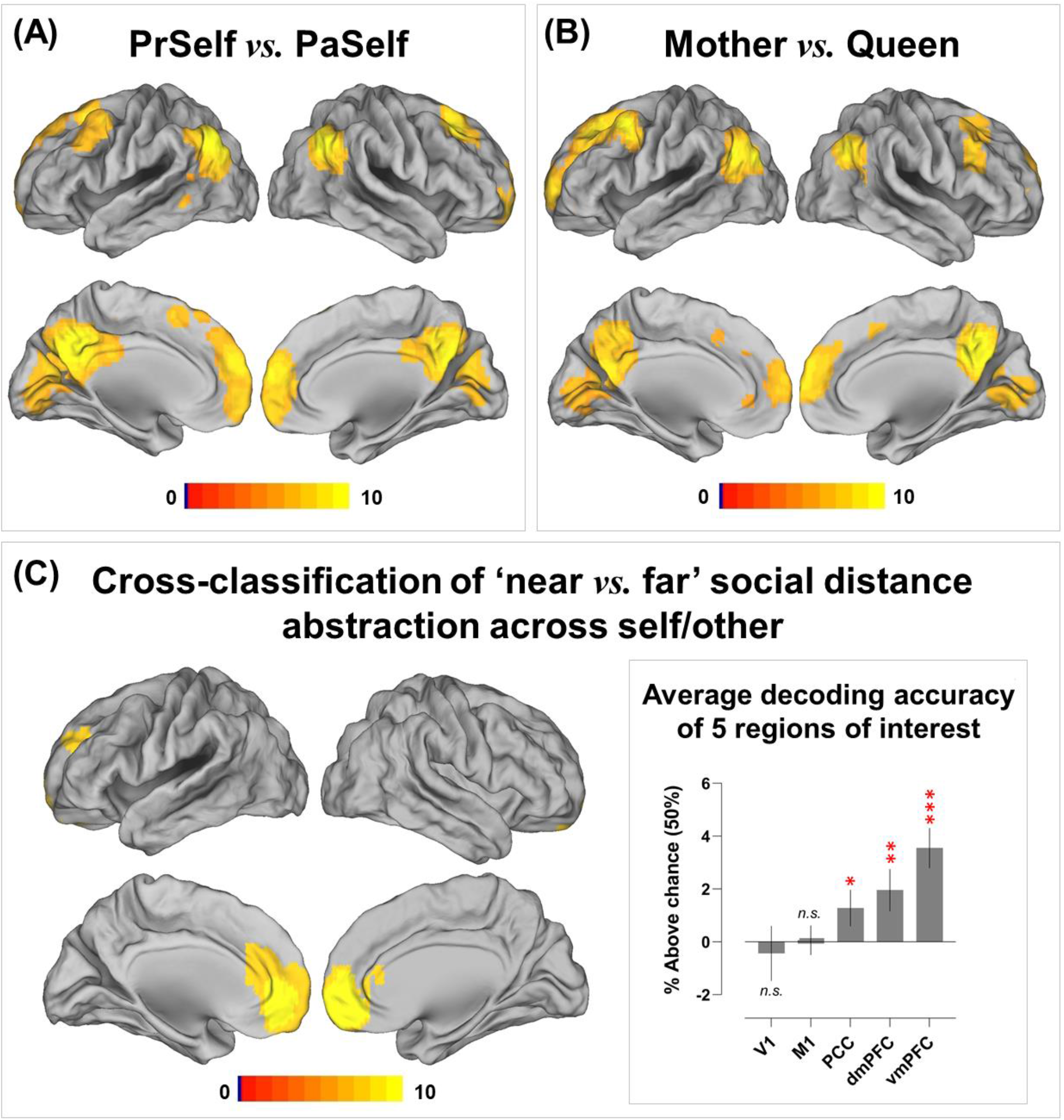
Whole-brain searchlight interrogation results for **(A)** Present Self *vs.* Past Self, **(B)** Mother *vs.* Queen, and **(C)** Cross-classification of ‘near *vs.* far’ social distance across the self and other domain. Shown in the inset-box is the extracted decoding accuracy from five target anatomically-defined ROIs (two control areas: V1/the primary visual cortex, M1/the primary motor cortex; 3 midline structures of the default- mode system: the PCC, dmPFC, vmPFC; these ROIs were selected *a priori* and defined using the anatomical masks of the Wake Forest PickAtlas Toolbox). Correction for multiple comparisons was via constraining voxel-wise FWE under *p* < 0.05

While the standard cross-validation approach unravelled whether the information of self *vs.* other existed in the multivoxel pattern of an ROI (e.g., whether the classifier succeeded in predicting if one was reflecting on *Mother* or *Queen* based on distributed activities), it lacked the ability to test whether the brain reinstates a reproducible and generalisable neural code across varying situations. A generalisable code indicates *abstraction* despite contextual variation. A hypothetical example that epitomises this generalisability and invariance would be that the brain summons reproducible neural patterns to represent one’s identity no matter whether that person is seen, heard, or recalled. To overcome the limitation of standard cross-validation, we conducted cross-classification – the classifier was trained on data from one cognitive context and tested on another; with this procedure, we tested whether there was any commonality in neural codes invariant to contextual changes. Two analyses were performed: First, we tested whether there is generalisable neural coding of self *vs.* other, irrespective of social distance. This was achieved by training the algorithm to classify *Present Self* from *Mother* (closer distance) and testing whether the codes were transferrable to distinguish *Past Self* from *Queen* (farther distance), and *vice versa*. Second, we tested whether generalisable/reproducible patterns of neural activities were used to represent the four individuals, irrespective of the emotive valence of descriptions that were used to probe person-related concepts. By virtue of our factorial design that crossed four identities with valence, this cross-classification was achieved by training the algorithm to classify the four persons using data from the positive-valence context and testing the generalisability using the negative-valence data (and *vice versa*). Cross-classification between contexts provides a rigorous test to assay whether invariant/generalisable neural codes were reliably summoned to represent the mental concepts about particular individuals, unaltered by social distance or emotive valence.

As illustrated in Figure 3(A), the analysis showed that, based on the neural patterns of four regions of the DN (the dmPFC, vmPFC, PCC, and left IPL), the classifier was able to extrapolate information (e.g., neural codes elicited by the pair of near distance – *Present Self vs. Mother*) from one context and successfully applied it to cross-classify in another context with a different degree of social distance (e.g., far: *Past Self vs. Queen*). This suggests context-invariant patterns that encode the *essence* of ‘self *vs.* other’, generalisable across close and distant social relationships. The representational content of these regions is instantiated in the three inset-boxes of Figure 2A. Here we saw a consistent pattern across the three ROIs with highest cross-classification accuracies - when the ground truth was ‘self’ the classifier was more inclined to predict ‘self’ than ‘other’, and *vice versa* when the ground truth was ‘other’. A coherent but more striking pattern was found in the four-way cross-classification which generalised across positive *vs.* negative emotive valence. As illustrated in Figure 3(B), based on the regional activity of an ROI, the algorithm was capable of extrapolating personal identities from one context and leveraging the codes to make predictions in another context of different emotive valence. Significantly above-chance cross-classification was achieved in every ROI of the DN and SN. The representational contents of such cross-valance neural coding were exemplified by the pattern of PCC activity: As illustrated in the inset box of Figure 3(B), the percentage of the classifier’s prediction clearly followed a sequence-like pattern (particularly conspicuous in the polar ‘extreme’ cases of *Present Self* and *Queen*). For instance, while the classifier correctly predicted *Present Self* most frequently when the truth was indeed *Present Self*, its erroneous responses conformed to social distance – *Present Self* was most confused with *Past Self*, followed by *Mother*, and least confused with *Queen*. A similar pattern (with the opposite order) was observed for *Queen*. Together, these indicated that the underlying neural representations that were employed to represent the distinction amongst personal identities were reproducible across contexts, invariant to changes of social distance and emotive valence, and were structured in a gradational manner that followed interpersonal distance.

**Fig. 3.**
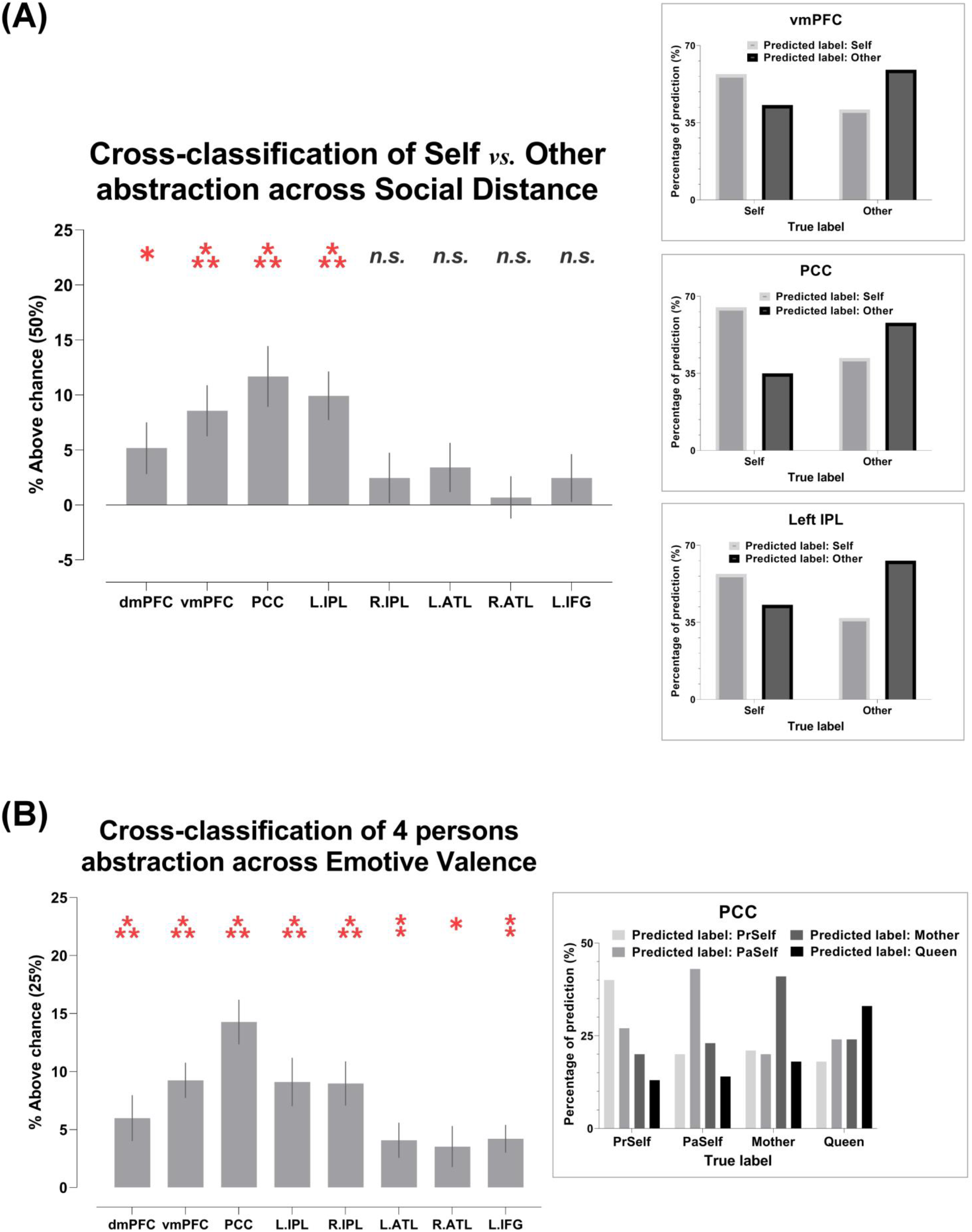
**(A)** Cross-classification of self *vs.* other between near and far interpersonal distances. **(B)** Cross- classification of four individual identities between positive- and negative-valence stimuli. Inset boxes show the confusion pattern of ROIs with highest cross-classification accuracy. Bonferroni-correction was applied for constraining multiple comparisons. **\*** *p* < 0.01; **\*\*** *p* < 0.005; **\*\*\*** *p* < 0.001

By visual inspection on cross-validation and cross-classification analyses, we noticed a disparity that decoding accuracy was obviously better in DN regions than in SN regions. This implies that DN regions might contain more person-related information than SN regions. To further investigate this gap in predictive power between the two subnetworks, we employed the combinatorial-ROI decoding analysis (for precedents, see Clithero *et al*. 2009; Smith *et al*. 2013; Wang *et al*. 2017). This analysis entailed a stepwise procedure wherein decoding was conducted on a ‘combined’ ROI - each combined ROI incorporated the pattern of an original ROI *plus* another ROI, followed by decoding analysis on the combined pattern; this was iterated for every pairwise combination of the eight ROIs. The outcome of joint-ROI decoding was subsequently compared with the ‘baseline’ where the decoding was conducted using the pattern from the original ROI alone. This procedure revealed whether adding a given ROI improved or impaired the classifier’s performance through serially coupling this ROI with all other ROIs and assessing how the joint-decoding altered relative to baseline. As has been demonstrated by previous research (Clithero *et al*. 2009; Smith *et al*. 2013; Wang *et al*. 2017), this method bypassed the difficulty of directly comparing between ROIs on the amount of information they carried (i.e., in the serial joint-decoding, the number and composition of voxels in a pattern were exactly matched; this way, we were able to quantitatively evaluate if an ROI a reliable ‘contributor’ or ’beneficiary’ when it was coupled with another ROI). We found that (*i*) adding any of the five DN regions to the combinatorial decoding robustly boosted accuracy whereas adding any of the three SN regions had little impact on decoding; (*ii*) the three SN regions reliably benefited more from the addition of another ROI whereas the DN regions benefited less. Shown in Figure 4 are the example results of joint-decoding for ‘*Self vs. Other*’ (Figure 4A), ‘*Present Self vs. Past Self*’ (Figure 4B), and ‘*Mother vs. Queen*’ (Figure 4C). A highly reliable pattern was seen in these results (as well as in other joint-decoding). Dovetailing our earlier data, the joint-decoding revealed a robust ‘imbalance’ between the amount of information carried by the two subnetworks. Whereas DN regions carried more information about personal identities (making them ‘givers’ that reinforced classification accuracy), SN regions carried less information (making them reliable ‘takers’ in the combinatorial-ROI decoding).

**Figure 4.**
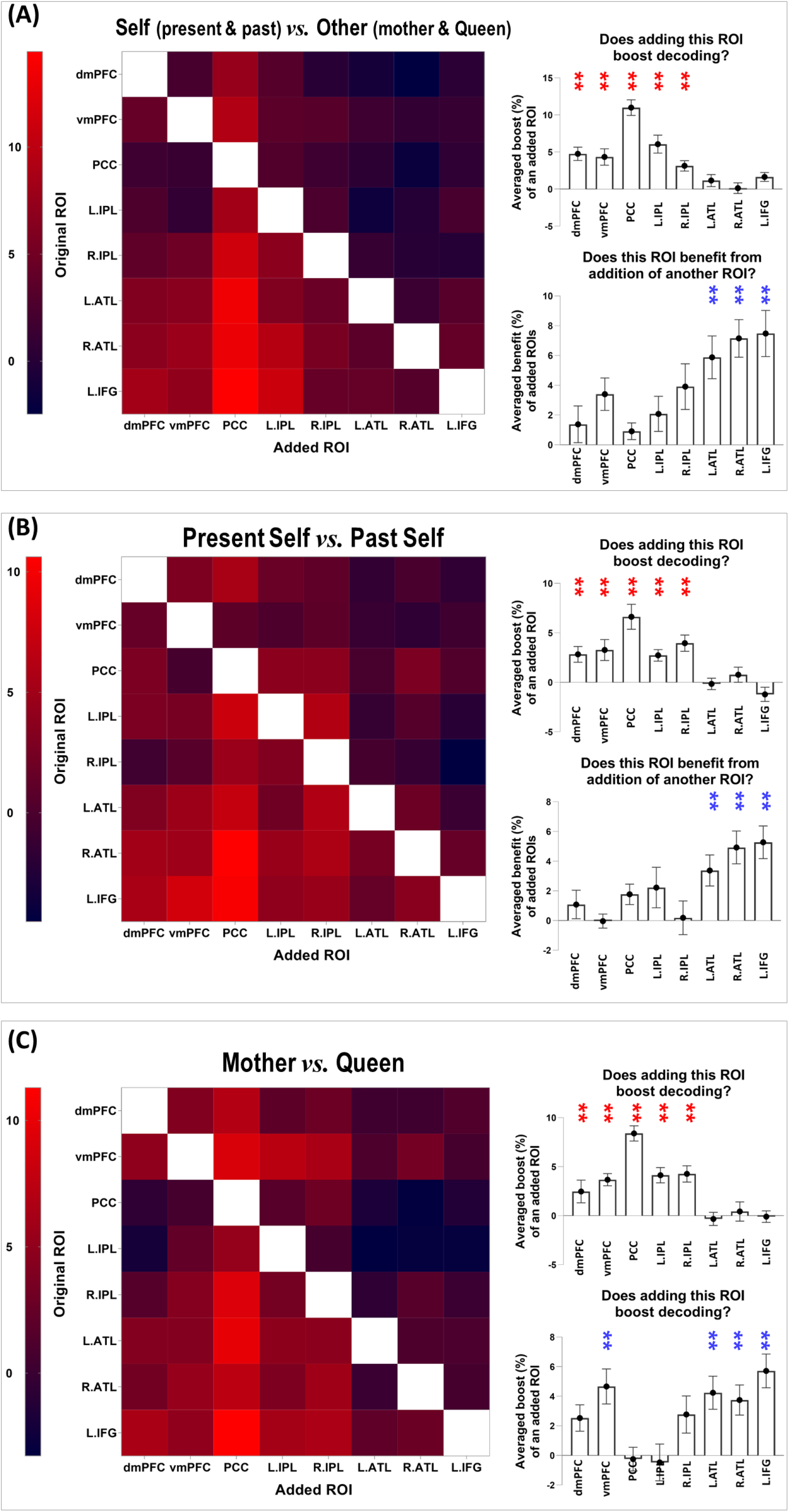
Combinatorial decoding that was based on an iteratively procedure that examined how decoding performance varied when the neural pattern of an original ROI was joined by the pattern of another ROI. The values on the colour scale indicates changes in decoding accuracy when an ROI was included, compared to the decoding result based on the original ROI alone, from dark purple (below zero, indicating *decrement*) to bright red (indicating most *improvement*). A reliable pattern was found across all analyses – as shown by the three examples here: **(A)** Present Self + Past Self *vs.* Mother + Queen; **(B)** Present Self *vs.* Past Self; **(C)** Mother *vs.* Queen. The bar-graphs indicate that, across analyses and across original ROIs, the addition of an ROI that belongs to the default network led to significant improvement of decoding results, whereas ROIs of the semantic network reliably benefited most from the addition of another ROI.

### Representational similarity analysis

Results of the classification analysis gave testable hypotheses about the configuration of neural coding: The brain might encode personal identities as continuous progression along a single-dimension spectrum of social distance, from close to distant, without any categorical cut-off. Alternatively, the neural codes could be configured with a boundary that categorically separates self-ness from otherness, with finer-grained differentiation nested within the ‘self’ or ‘other’ domain. The confusion matrices of classification analysis (see *Supplemental Results 6*) only provided an equivocal answer – although the patterns of confusion showed that within-domain confusion was more frequent than between-domain (e.g., *Queen* was more confused with *Mother* than with the two variants of self, which implied a bipartite structure), such analyses did not directly quantify the extent of the similarity between two representations. Unlike classification-style analysis that by nature discretises different categories via imposing a decision boundary, RSA quantifies the extent of similarity using continuous measures (e.g., correlation coefficient). Therefore, we exploited RSA to further investigate the representational ‘geometry’ that underlies the similarity between different categories and different brain regions.

We first verified whether there was good concordance between the neural representations of positive- and negative-valence conditions, given the fact that the SVM algorithm successfully cross-classified individual identities across valence. As illustrated in Figure 5, in every ROI, a statistically robust correlation was found between the representational matrices of positive- and negative-valence contexts, suggesting a coherent motif that the classifier could extrapolate from one situation and apply to another. Moreover, the outcomes of RSA and pairwise-classification concurred with one another. An example result from one ROI is illustrated in Figure 6: When the representational similarity was high between two conditions (which indicated a higher degree of overlap in neural coding), there was a corresponding decline in the accuracy scores of classification (the classifier was more prone to confusion). This resulted in negative correlations, reliably seen in both positive- and negative-valence contexts. Next, hierarchical clustering was used to visualise the representational distance between conditions, quantified by the branching of a dendrogram tree. As Figure 7 illustrates, in these four ‘core’ regions of the DN (in which we saw generally higher classification accuracy), there is a clearly dyadic structure that stratified neural representations into two major branches along the border of ‘self *vs.* other’. One branch comprised *Present Self* and *Past Self*, while the other branch comprised *Mother* and *Queen*. Interestingly, in the vmPFC, the representations of *Present Self* (encompassing both positive and negative) formed a distinct cluster that was separable from the remainder (even detached from *Past Self*). This is consistent with the literature that the vmPFC has a unique role in representing the ‘essence’ of self-concept. It is also clear that emotive valence has minimal impact on the segregation between neural representations, which further validates a reproducible coding of personal identities unvarying across valence. Whereas a clear bipartite split between self and other was found in the representations of DN regions, in the three SN regions we did not see any systematic clustering that demarcated the difference between conditions. Taken together, these results emphasise the convergence of evidence that we obtained from the RSA/correlation and SVM/classification approaches.

**Figure 5.**
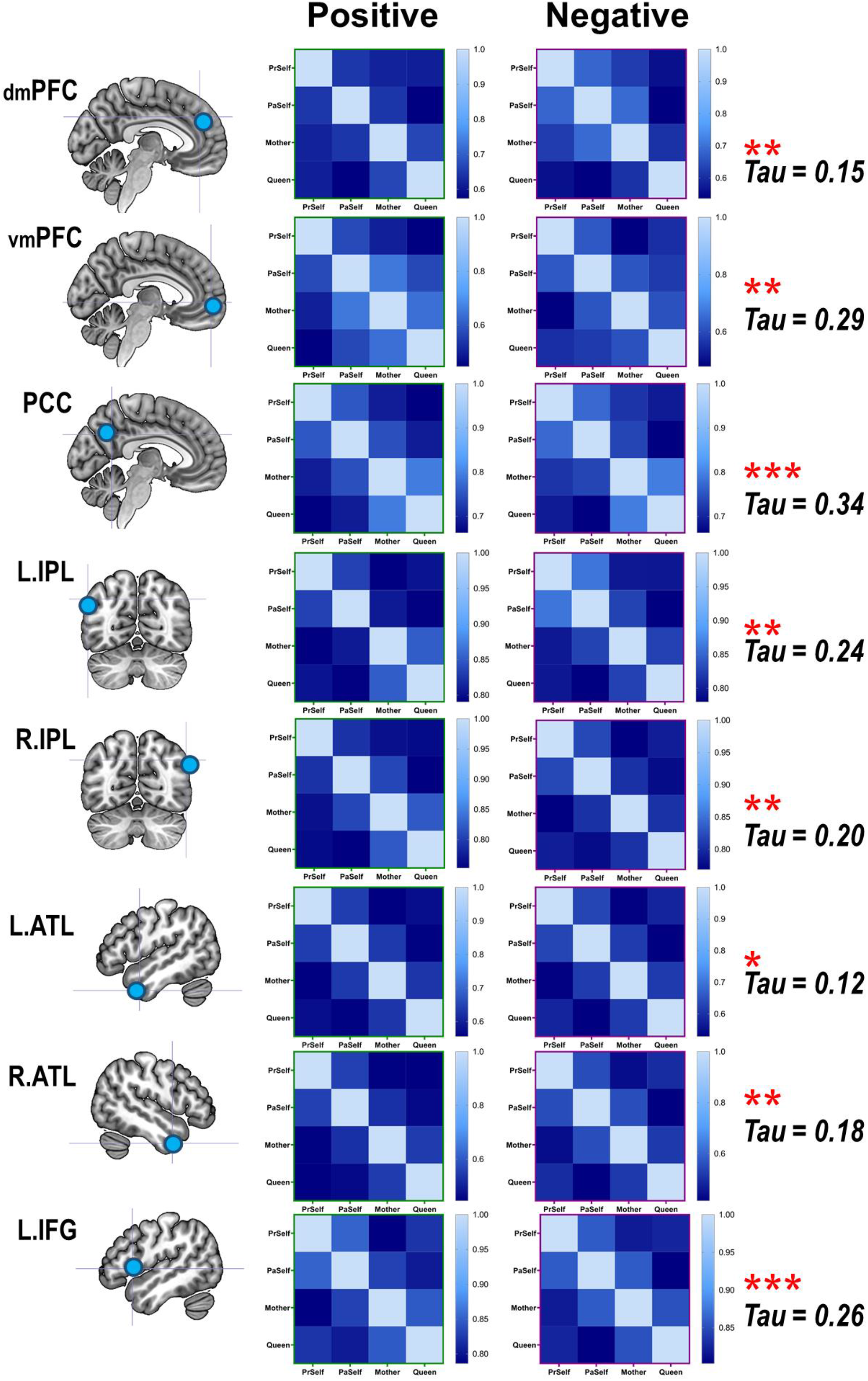
Corroborating the results of cross-classification between positive- and negative-valence stimuli, the results of representational similarity analysis showed that the neural patterns elicited by the four personal identities are significantly correlated between the contexts of positive- and negative-valence. The correlation was found in every ROI of the DN and SN.

**Figure 6.**
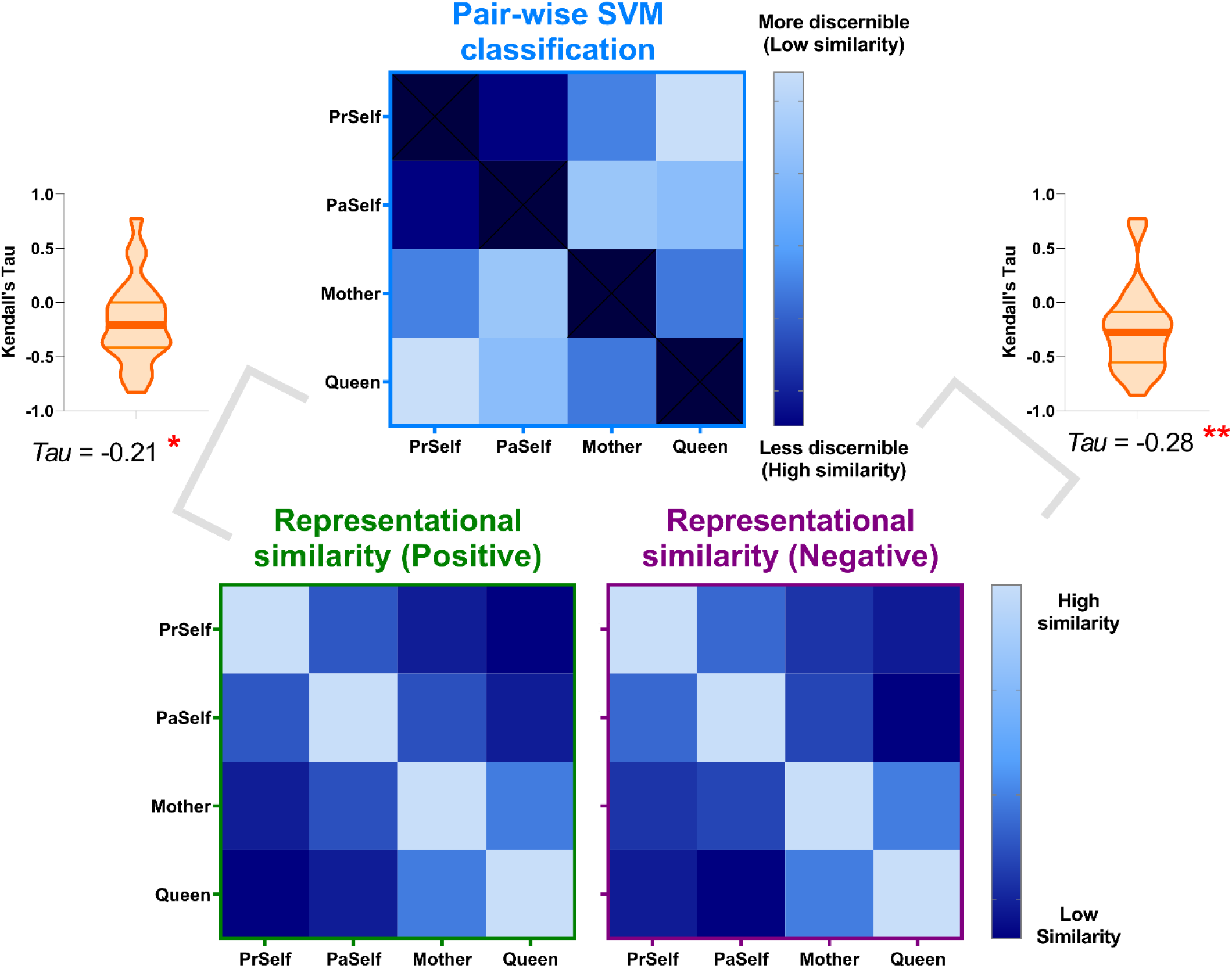
The results of representational similarity and machine-learning classification analyses dovetailed with each other. Illustrated here are example results based on the neural patterns of the posterior cingulate cortex (coherent results were observed for all other regions) – when two experimental conditions were representationally *more* similar to each other (hence more confusable), the classifier was *less* able to predict the correct category label, resulting in significant negative correlations found in both positive- and negative-valence conditions.

**Figure 7.**
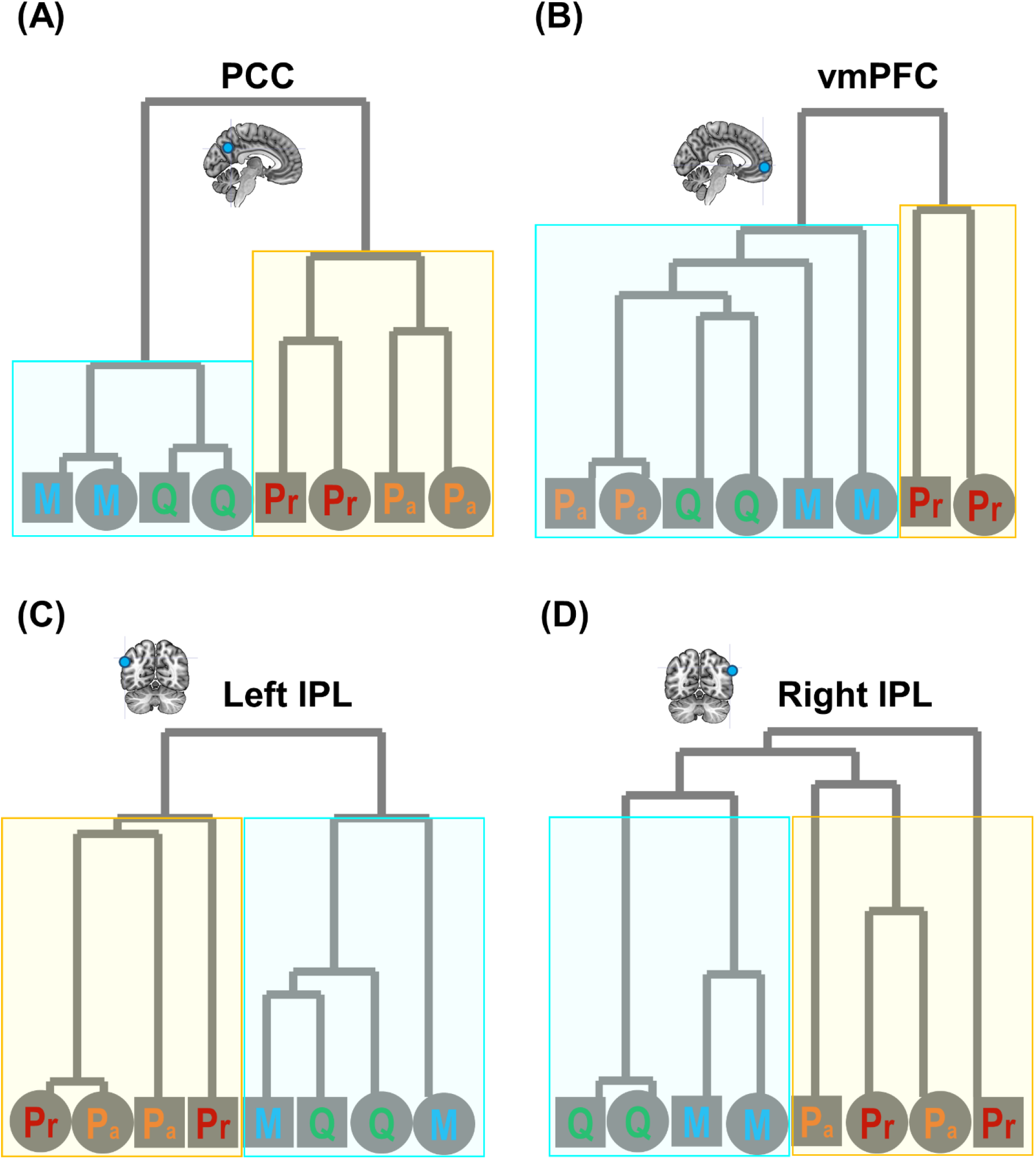
The configurations of each dendrogram tree here show a clear separation between an aggregate class of *self-ness* (*Present Self* and *Past Self*; colour-coded using red and orange, respectively) and aggregate sense of otherness (*Mother* and *Queen*; coded using blue and green, respectively). Corroborating the cross-valence classification, the structure of representational dissimilarity (1-*r*) showed that emotive valence had little impact on clustering arrangement (which indicates the extent of neural similarity). Social distance (coded using different colours) indeed impacted on the clustering, but it was couched within the ‘self/other’ separation. Emotive valence is coded by geometric shapes: circle – positive, square – negative. This reliable broad-brush segregation between self and other is found in core DN regions of **(A)** PCC, **(C)** left IPL, and **(D)** right IPL. The vmPFC – in Panel **(B)** – is somewhat different that it treats *Present Self* as a unique category different from all other categories.

Replicated across multiple classification analyses, decoding accuracy was found to be robustly higher in DN regions compared to SN regions. Furthermore, combinatorial-ROI decoding also showed that the addition of DN regions reliably boosted accuracy whereas adding SN regions caused no improvement, suggesting an asymmetry of information quantity between the networks. To investigate this issue further, we used RSA to examine whether this asymmetry translates into representational distance between the two networks. Hierarchical clustering showed that the principal factor that characterised the heterogeneity between neural representations was the separation between DN and SN. As the configuration of the dendrogram illustrates (Figure 8A), the initial bifurcation between regions was driven by the network membership an area belongs to. Within the cluster of DN, there were two sub-clusters – the two medial prefrontal regions (the dmPFC and vmPFC) formed a subgroup separable from the posterior regions (medial-parietal: the PCC; lateral-parietal: the left/right IPL). Within the cluster of SN, the clustering was consistent with contemporary theories and findings of semantic cognition (e.g., Lambon Ralph *et al*. 2017) – the areas representing semantic meaning *per se* (the left/right ATL) formed a sub-cluster separable from the area that controls the retrieval of semantic meaning (the IFG). This structure is also evident in the similarity matrices whereby we colour-coded the strength of correlation/similarity (Figure 8B): Close inspection of the layout of correlation matrix revealed that the three SN regions congregated to form a cluster that was separable from all of the remaining DN regions; within the five DN regions, the dmPFC and vmPFC formed a subcluster while other posterior regions formed another subcluster. These two visualisation methods provide complementary yet consistent insight into the comparison between neural networks. Finally, we computed the averaged sizes of correlation between brain regions as a function of whether it was correlating two regions within the same network or between the two networks, as well as whether data was based on the positive- or negative-valence context. Results showed a significant effect of network membership – correlation size was significantly greater within the same than between networks (Kendall’s τ-a: *F*_(1,23)_ = 6.36, *p* = 0.01, *η*_p_^2^ = 0.22), indicating greater representational similarity between regions of the same clan. By contrast, valence had no effect (again indicating a coherent representational structure across valence), nor did the valence × network interaction (both *ps* > 0.55). To ascertain robustness, we repeated all these analyses using a different type of rank-correlation measure (Spearman’s *ρ*) and obtained entirely consistent results (see Figure 8). Together, these results complement our observations of the classification-based analysis and further highlight the subtlety of subdivision between networks (and within a network).

**Figure 8.**
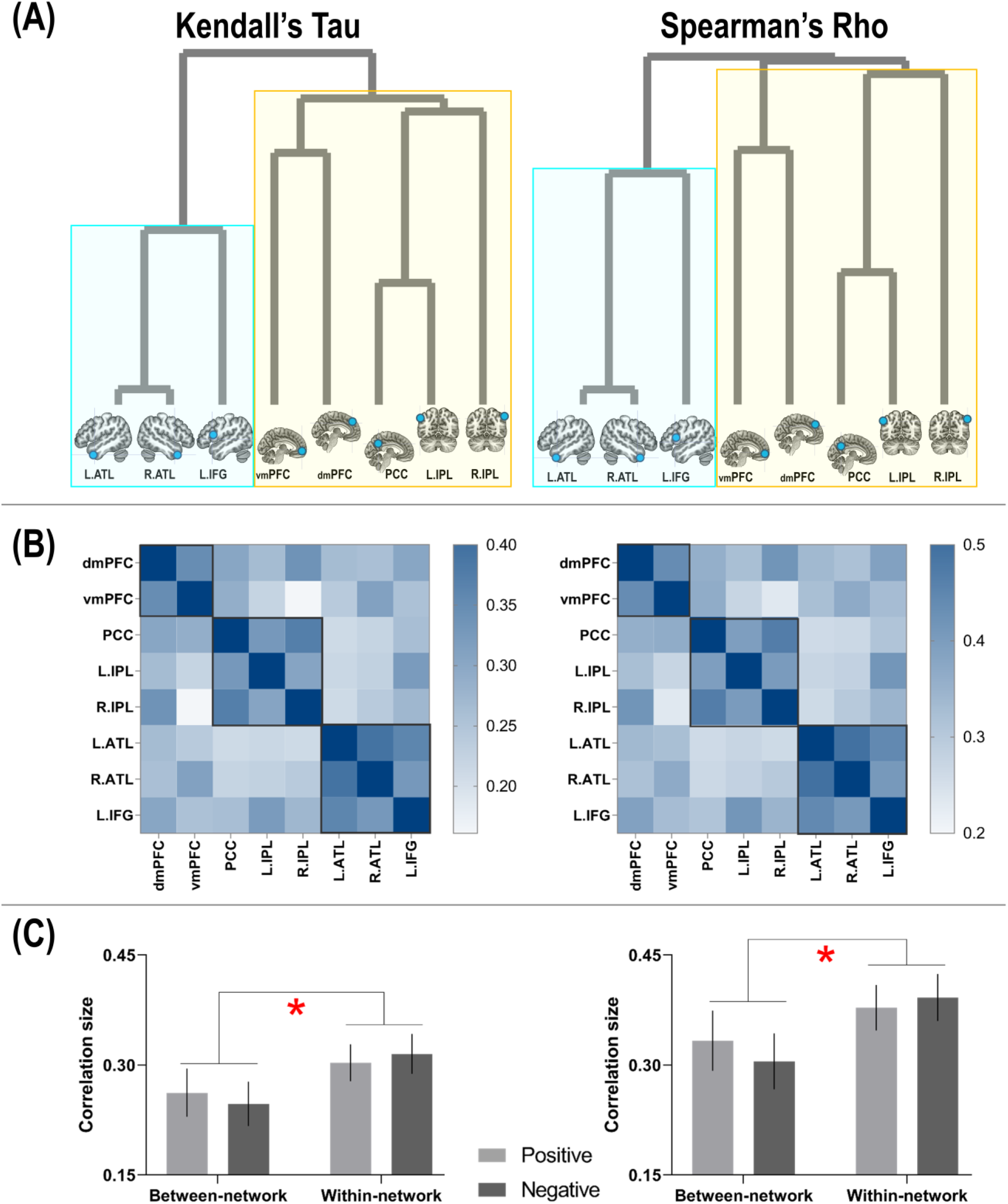
**(A)** The configurations of dendrogram show that representational distance (1 minus τ-a or *ρ*) dissociated between the semantic network and default-mode network; this was seen both using Kendall’s Tau-a and Spearman’s Rho; **(B)** Here representational similarity is delineated using correlation matrix. A consistent pattern with the dendrogram structure is highlighted using black boxes; **(C)** Within-network representational similarity is significantly higher than between-network similarity, reliably found in both the contexts of positive- and negative-valence.

Finally, we examined three theoretical models, testing their explanatory power to account for the neural data (see Figure 9). The first model was derived from each participant’s subjective rating of the pairwise similarity between the personality traits of two individuals using a continuous scale (most dissimilar: 0 – most similar: 100). The second model was a hypothetical model based on a binary distinction between self and other – *Present Self* and *Past Self* were collapsed under the ‘self’ umbrella while *Mother* and *Queen* were under the ‘other’ umbrella. The third model was a hypothetical model based on gradual changes, with each of the four individuals assumed to be equidistant from one another on the spectrum. We correlated the neural similarity matrix of each brain region with these theoretical models, and statistically examined its reliability (FDR-adjusted for multiple comparisons). We found that the neural pattern of every brain region significantly correlated with the binary model and the graded model (all *p*s < 0.005). As illustrated in Figure 9, for both of the binary and graded model, the neural pattern of the PCC was most correlated with the theoretical models and approached the lower-bound of noise ceiling (indicating the models’ near-optimal capability to explain neural data). The correlation sizes with the theoretical model did not reliably differ between brain regions, nor did the comparison between the binary model and graded model (all *p*s > 0.05 with FDR-correction). By contrast, the behavioural rating matrix did not reliably correlate with any brain region. Critically, both the binary and graded models outstripped the model of behavioural rating in terms of their explanatory power on the neural data (binary *vs.* behavioural: *p* = 0.0008; graded *vs.* behavioural: *p* = 0.001). While subjective rating best characterised how a participant perceived and compared the resemblance of two characters, such high-dimensional, multi-faceted understanding (quantified as their rating) was not fully reflected in the local neural patterns of DN and SN regions. Instead, while the binary model was less elaborated (or more impoverished in the dimensions to characterise the difference of personality) than behavioural rating, it was better able to capture the ‘representational landscape’ of neural data. This concurs with earlier results that the broad-brush separation of ‘self *vs.* other’ explained most variance. By contrast, while socially ‘close *vs.* distant’ was decodable from the cross-classification between self and other (implying that it could be an orthogonal dimension to self/other), it was a less dominant factor in shaping the landscape of neural representations.

**Figure 9.**
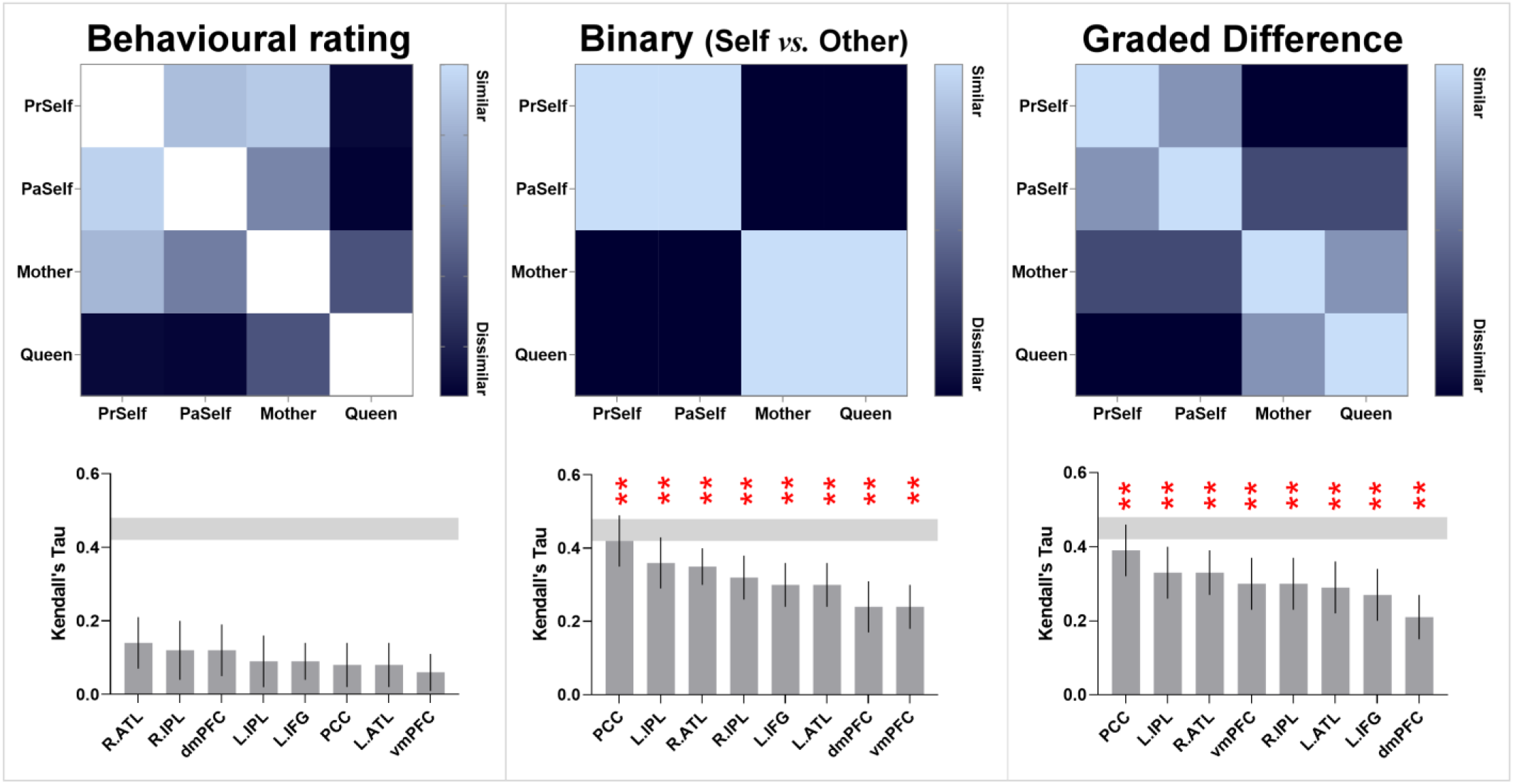
The hypothetical models that assumed a binary difference between self and other (middle) and gradational differences of the four individuals (right) are significantly correlated with the neural pattern of every region. The noise-ceiling is averaged across all regions. Multiple comparisons are controlled by constraining FDR (*q*) under 0.05. *\*\* p* < 0.005. By contrast, the outcomes of behavioural rating (left) were not correlated with the neural pattern of any region.

## Discussion

In the present study, we used a series of multivoxel decoding to unravel the neural representations of self- *vs.* other-referential concepts, a pivotal psychological construct that permeates all aspects of social life and has been argued as a core function of the specialised system for social cognition. Across multiple analyses, we found a robust dyadic fractionation between the two subnetworks within the social system – multivoxel patterns reliably dissociated between regions that are canonically affiliated with the DN and those with the SN, evident both in the outcomes of supervised classification and representational similarity. Moreover, representational configuration of the neural response to self- and other-referential thoughts mirrored the psychological continuum of interpersonal distance, from an integral sense of *Present Self* to a distant other (*Queen*), with the broad-stroke segregation of ‘self-ness’ from otherness’ being the chief factor that divided the representational space, and social distance^2^ being an auxiliary factor that subdivided concepts within the self/other domain. Below we discuss the implications of the present results.

### The fusion and fission of DN and SN

In the literature of social neuroscience, various regions of the DN and SN have been demonstrated to represent diverse mental states (e.g., Tamir *et al*. 2016) and personality traits (e.g., Thornton and Mitchell 2018), which leads to an agglomeration of the two networks as the social system. In two separate bodies of literatures, however, neurolinguistics and human connectome research have demonstrated that the DN and SN are two entities with distinct functional/anatomical profiles but they unite and disjoin in various resting- and task-states. Data-driven parcellation methods have demonstrated that DN and SN regions form a coherent network at rest (e.g., Yeo *et al*. 2011). Seed-based connectivity methods have also shown that key nodes of the SN – the ATL and IFG – are connected with DN nodes at rest (Jackson *et al*. 2016; Humphreys and Lambon Ralph 2017). Their association during resting-state has also been found during task-states: DN and SN activities are both enhanced by mnemonic representations (e.g., autobiographical memory) and inhibited by externally-driven processes (sensory input or motoric output; Chiou *et al*., 2020). However, the two networks have also been found to reliably dissociate: Chiou *et al*. (2020) demonstrated that, while both networks showed heightened activation for socio-cognitive tasks, DN regions preferred tasks emphasising the retrieval of episodic details whereas SN regions preferred tasks emphasising the semantic interpretation of perceptual input. Jackson *et al*. (2019) and Humphreys *et al*. (2015) both found that typical semantic tasks drove a dissociation – they activated SN regions but suppressed DN regions. Given the mixed picture, the findings reported in the present study have important implications for clarifying the relationship between SN and DN – we demonstrated that both networks contained information for decoding social categories; however, compared to the SN, the DN possessed more abstract/generalisable information that allowed cross-classifying across distance and valence, and exhibited sharper representational partitions that individualised each social category. The disparity between the two networks has implications for constraining the interpretation on the activity of SN regions during various socio-cognitive tasks (e.g., Olson *et al*. 2013; Binney *et al*. 2016; Wang *et al*. 2017) – for instance, if some SN regions prefer social knowledge to other semantic contents, there might be more exchange of information between these SN regions and DN regions that could be unravelled using connectivity analysis, compared to SN regions that prefer non-social contents. More broadly, human connectome research has shown that both the DN and SN are situated at the high-order, transmodal end of macroscale cortical hierarchy (Margulies *et al*. 2016). In this regard, our finding may pave the way for future research to clarify how their positions in this macroscale architecture affects the information they carry. We elaborate on this in the following section.

### The bipartite split in multivoxel patterns between networks

Decades of research have accumulated a wealth of data about how inter-connected brain regions form large-scale networks, as well as how network architecture can be mapped onto cognitive functions (for review, Uddin *et al*. 2019). Of particular interest is the functionality of the expansive default-mode network. This widely dispersed group of brain areas were originally identified based on their ‘deactivation’ (relative to task-state) during active engagement in a task-state and heightened activation-level during wakeful resting periods (Raichle *et al*. 2001). Owing to its initial ‘task-negative’ definition, for a long time this system was assumed to play little role in goal-oriented behaviour. Later research has identified its contribution in a panoply of goal-directed cognitive tasks that depend on internally-constructed representations (e.g., memory, schema, etc.), such as social cognition (e.g., Skerry and Saxe 2015), retrieval of episodic/autobiographical memory (e.g., Spreng and Grady 2010), skilful application of schema to solve a task (Vatansever *et al*. 2017), and short-term visual memory (as compared to visual perception; see Murphy, Wang*, et al.* 2019). Despite mounting evidence, the ‘misnomer’ of default-mode system (rooted in its original task-negative definition as the brain’s metabolic default) continues to be used extensively. Recent research has reliably uncovered a bipartite structure within the default system (e.g., Braga and Buckner 2017; Braga *et al*. 2019; Chiou *et al*. 2020; DiNicola *et al*. 2020; also see Andrews-Hanna *et al*. 2010 for a tripartite fractionation of the DN). The bipartite structure has been discussed under the framework of a macroscale ‘gradient’ spanning the entirety of cerebrum (Margulies *et al*. 2016; Huntenburg *et al*. 2017). This gradient reflects a recursive process of information convergence, occurring in multiple places of the brain, from sensory-motoric representations that encode the ‘here-and-now’ of perceptible entities to multisensory representations that are stored in the DN/SN and encode memories and concepts. Core regions of the DN (e.g., the PCC and vmPFC) sit atop this gradient, while core regions of the SN (e.g., the ATL and IFG) are situated at a tier below the apex position (i.e., the DN’s cores). The dissociation between DN and SN might reflect their differential position on this gradient of information convergence. Our present multivoxel-decoding results lend support to this framework – as revealed by both classification- and similarity-based analysis, we found that DN regions contained more information about individual identity, which is an abstract concept distilled from multiple sources of sensori-motoric modalities and accumulated over multiple life-events. Thus, compared to SN regions, the more informative DN regions enabled better classification accuracy (as evident in the outcomes of iterative joint- decoding analysis) and exhibited more clear-cut representational segregation between self and other (as evident in the branch of dendrogram trees). These findings are compatible with the interpretation that the DN contains more abstract information for social cognition, despite the fact that the DN and SN both enhanced their activation in response to the demand of social processing (in a univariate ‘BOLD amplitude’ sense).

Significantly above-chance decoding was found in both networks for coarser- and finer-grained classifications of self *vs.* other, indicating that both of them carried sufficient information that allowed the algorithm to delimit a margin to distinguish between two/four classes of data-points. Amongst all ROIs, the PCC contained most information about personal identities. This is observed across multiple analyses – decoding accuracy was highest based on the patterns of PCC; when the PCC was added to the combinatorial decoding, it led to greatest boost the outcome of decoding; the neural pattern of PCC was most correlated with the psychological continuity of self *vs.* other, leading to correlations that approached the noise ceiling (indicating near optimum). These data are consistent with the view that the PCC serves as one of the cores of the DN (Andrews-Hanna *et al*. 2010) such that its representational pattern ‘echoes’ the activities elsewhere in the brain through long-range connections (Leech *et al*. 2012) and its dysfunction causes memory-related ailments (Leech and Sharp 2014). Moreover, it is noteworthy that the vmPFC represents *Present Self* as a distinct entity from all other categories, as evident in how the dendrogram initially bifurcates in Figure 7B. This result is consistent with the specialised role of the vmPFC in representing the essence of self-concept, adding to a large body of evidence (for review, Wagner *et al*. 2019). Recently, the ‘self-in-context’ model about vmPFC function has been proposed (Koban *et al*. 2021) – according to this view, the vmPFC represents ‘self’ in a compressed low-dimensional space that captures relevant features of a context (e.g., human interaction, social norm) to construct ‘self’.

It is important to note that any individual brain region may participate in multiple brain networks and subserve multiple cognitive functions even though a region has a primary network affiliation or has certain types of functions that it is frequently associated with (Pessoa 2014). The ‘network membership’ of a brain region can be affected by various factors, such as its position in the network (i.e., ‘connector’ regions between two networks tend to have more fluid network affiliation, relative to regions within the ‘heartland’ of a module; see Spreng *et al*. 2010; Spreng *et al*. 2013), the cognitive state one is under (i.e., contexts can drive a region to fluidly couple with different networks), as well as the methods used to probe the relationship between regions (for discussion, see Petersen and Sporns 2015). These factors may particularly affect several ‘connector’ regions that bridges between two networks, making them exhibit characteristics of both systems (such as the dmPFC that bridges core DN nodes with SN nodes; see Spreng *et al*. 2013). We speculate that this may offer explanations regarding the fluid functional profile of dmPFC: Previously, using *univariate* analysis, we found that the dmPFC responded preferentially to semantic tasks over episodic tasks, which makes it more akin to the behaviour of SN regions (e.g., the IFG and ATL) than DN regions (e.g., vmPFC; Chiou *et al*., 2020). However, using *multivariate* analysis, in the present study we found that representational contents of the dmPFC was more similar to those of DN regions relative to SN regions, making it more affiliated to the DN. Together, our data are consistent with the previous literature concerning this region’s somewhat inconclusive affiliation with different networks; it also highlights the flexibility of a ‘connector’ region and the peril of imposing a rigid demarcation on the perimeters between networks.

### The representational structure of self- vs. other-concept

Our findings add to the literature that deciphering self-/other-referential thoughts is possible based on the patterns of DN and SN regions (Hassabis *et al*. 2013; Thornton and Mitchell 2017; Courtney and Meyer 2020; Peer *et al*. 2021). Moreover, our cross-classification analyses have important implications for the cognitive theories of self-processing. The ability to successfully cross-classify has been considered as a benchmark of testing whether there is genuine abstraction of neural coding across domains (Kaplan *et al*. 2015). We found that the brain uses robustly contextually generalisable neural patterns to encode self-ness and otherness such that this cardinal representational code applies across near and far interpersonal distances and across positive- and negative-valence depictions. On top of the separation between self and other, social distance was another factor that sculpted the representational landscape of the neural codes. As discussed earlier, ‘self vs. other’ serves as the primary dimension that delineates the most salient difference between classes, while other auxiliary dimensions (e.g., social distance) are couched within the primary segregation. This is reminiscent of representational dimensions of visual perception (Konkle and Caramazza 2013): Animacy (living *vs.* non-living) is the primary dimension that separates animate entities from inanimate items, while object size serves as the secondary dimension that assorts subgroup within the inanimate class (Long *et al*. 2018). Interestingly, participants’ behavioural rating on personality was not correlated with the neural pattern of any region. A possible reason of his result might be that while the local pattern of an ROI is sufficient to represent (***i***) the binary/broad-stroke difference of self *vs.* other and (*ii*) the extent of social distance that intersects with the ‘self/other’ dimension, it was insufficient to capture the more sophisticated pattern inherent in the thoughts behind the behavioural rating. As the matrix of behavioural rating illustrates (Figure 9), our participants rated that their own mother was more similar to themselves and drastically different from the *Queen*; this led to a clear cluster that lumped the three personally familiar targets together (*Present Self*, *Past Self*, and *Mother*), separate from the *Queen*. The majority of participants rated that the similarity of *Present Self* and *Mother* was even higher than that of *Present Self* and *Past Self*, which reflects complex, metacognitive thoughts that apparently transcend the ‘self *vs.* other’ boundary. This implies elaborated and multifaceted considerations behind the ratings that might require additional neurocomputation beyond the information within a single DN/SN region, whose local pattern cares primarily about ‘self *vs.* other’ *plus* social distance.

## Conclusion

In the present study, we reported evidence of multivoxel decoding that manifested a robust bipartite split within the brain’s social system into the DN and SN. The multivariate findings extend beyond previous evidence that was based on connectivity by demonstrating that different subnetworks within the social system dissociated on the amount of information that a subnetwork was laden with, as well as on the representational geometry that encoded person-related concepts. These findings inform the burgeoning field of human connectome research and its relationship with social neuroscience; our results also shed light on the decade-long investigation into how the brain implements the conceptual distinction of ‘self *vs.* other’.

***^1^***Here we used a well-established localiser paradigm (Dodell-Feder *et al*., 2011) to identify ROIs, pinpointing the peaks in the DN and SN that had greater activation for the Social than Non-social condition. To ascertain the robustness of our findings, we conducted three additional analyses: **First**, we used the coordinates from activation-likelihood estimation (ALE) meta-analyses on the literatures of semantic processing and social cognition to re-localise ROIs, and performed multivoxel decoding at these literature-defined locations (*Supplemental Results 2*). The decoding results based on ALE are highly consistent with those based on the Dodell-Feder localiser, demonstrating the robustness of our findings that they are not reliant on a specific localiser paradigm. **Second**, we used NeuroSynth to identify the voxels robustly engaged by semantic tasks (based on 40,030 activations from 1,031 fMRI studies; *Supplemental Results 3*). We found that all of the three semantic ROIs defined by our localiser contrast are situated within the semantic clusters of NeuroSynth, suggesting concordance between our semantic ROIs and previous neurolinguistics literature. **Third**, we used NeuroSynth to identify the voxels robustly engaged by self-related processing (based on 4,728 activations from 166 neuroimaging studies; *Supplemental Results 4*). We found that all of the five default-mode ROIs defined by our localiser contrast are situated within the self-related clusters of NeuroSynth, suggesting agreement between our default-mode ROIs and previous literature of self-processing.

^2^ It is noteworthy that social distance, as an auxiliary factor, was one of a multitude of factors that could determine the partition of social categories. For instance, in the differentiation of Present Self *vs.* Past Self, temporal distance was a factor that was closely entangled with social distance and could be an alternative explanation that configured neural representations.

## Acknowledgements

This research was funded by a Sir Henry Wellcome Fellowship (201381/Z/16/Z) to RC, and an MRC programme grant and intramural funding to MALR (MR/R023883/1; MC_UU_00005/18).

