## Supplemental Information for "Bipartite functional fractionation within the neural system for social cognition supports the psychological continuity of self *vs.* other"

Rocco Chiou[1] ✉, Christopher R. Cox[2], & Matthew A. Lambon Ralph[1] ✉
[1] MRC Cognition & Brain Sciences Unit, University of Cambridge, UK
[2] Department of Psychology, Louisiana State University, USA

✉ Correspondence to this work can be addressed to either of the authors by email:
(**RC:**, **MALR:**)

**Supplemental Stimuli:** Text presented in the task of personality assessment

| Positive valence | Negative valence |
|---|---|
| Open to new ideas | Anxious about uncertainty |
| Stoic to adversity | Patronising in attitude |
| Quick to understand | Aloof to people |
| Careful when preparing | Hesitant in taking chances |
| Independent in daily life | Cynical and sceptical |
| Charitable to people | Dishonest in behaviour |
| Articulate when speaking | Extravagant when spending |
| Easy to deal with | Jealous of others |
| Forgiving of others' mistakes | Impatient while waiting |
| Optimistic about future | Impulsive when acting |
| Creative and talented | Gullible and naïve |
| Confident in self | Complacent in attitude |
| Fond of joking | Distressed too easily |
| Humble and self-effacing | Fond of gossiping |
| Mature and steady | Selfish and uncaring |
| Mild in temper | Ashamed about mistakes |
| Graceful in manner | Boastful when speaking |
| Compassionate to others | Fond of binge drinking |
| Willing to adventure | Feeling incompetent |
| Generous with money | Snobbish in attitude |
| Sincere to friends | Having hypocritical behaviour |
| Diligent at work/school | Self-conscious when speaking |
| Able to be trusted | Bitter about the past |
| Efficient in planning | Aggressive when arguing |
| Candid during communication | Unrealistic when planning |
| Adaptable to new situations | Aimless about future |
| Insightful when making decisions | Insecure in relationship |
| Proficient in learning languages | Regretful about the past |
| Cheerful in disposition | Worried about trivial things |
| Keen to help | Sensitive to criticisms |
| Able to multi-task | Demanding and controlling |
| Determined to accomplish | Unpredictable in temper |
| Neat in housekeeping | Low in self-esteem |
| Punctual at work/school | Too eager to show off |
| Proactive in action | Volatile in mood |
| Diplomatic when talking | Too harsh when commenting |
| Organised in planning | Forgetful in daily life |
| Experienced in travelling | Overly suspicious |
| Poetic in writing | Showy and attention-seeking |
| Popular with people | Overconfident about ability |
| Having great charisma | Too lazy to tidy up |
| Hospitable to guests | Pessimistic about future |
| Carefree in life | Stubborn in thinking |
| Courteous to people | Reckless and careless |

| Positive valence | Negative valence |
|---|---|
| Relaxed in mood | Opinionated in communication |
| Athletic and energetic | Too fearful of failure |
| Courageous in action | Overly frugal with money |
| Motivated to succeed | Unwilling to open up |
| Prudent when spending | Clumsy and awkward |
| Outgoing and approachable | Arrogant in attitude |
| Chatty and humorous | Unforgiving to people |
| Able to stay resilient | Condescending in manner |
| Pragmatic in problem solving | Childish and frivolous |
| Patient while waiting | Evasive and dodgy |
| Laid-back in manner | Withdrawn in social contexts |
| Collegial in teamwork | Narcissistic and egocentric |
| Satisfied with life | Rigid in thinking |
| Genuine in attitude | Unhappy about life |
| Caring to people | Fickle when planning |
| Able to keep promise | Sloppy in writing |
| Cordial in manner | Complaining too much |
| Perceptive in spotting errors | Feeble and ill |
| Eloquent when persuading | Timid in temper |
| Unassuming in manner | Passive in action |
| Even-tempered and calm | Too clingy to friends |
| Devoted to family | Restless under stress |
| Respected by others | All talk no action |
| Earnest in attitude | Dull and unsociable |
| Dedicated to goals | Too fussy when choosing |
| Flexible in thinking | Insincere in attitude |
| Agile in movement | Merciless in attitude |
| Tolerant to different views | Overly self-critical |
| Rational in thinking | Cunning in planning |
| Assertive when speaking | Unimaginative in thinking |
| Liberal in attitude | Unkind when speaking |
| Encouraging to people | Disheartened about life |
| Upright in character | Quarrelsome and irritable |
| Sympathetic to people | Estranged from family |
| Keen to learn new skills | Resentful about others |
| Well-read and cultured | Prejudiced in attitude |

**Supplemental Results 1:** Behavioural data and decoding at the primary motor area

As illustrated in the figure below, assessing personality traits of the four target individuals results in comparable reaction times for the four conditions. Statistics ($F_{(3,69)} = 2.95$, $p = 0.04$) showed that no post-hoc pair-wise comparisons revealed any reliable difference excpet for reacting to *Present Self* being significantly quicker than reacting to *Past Self* ($p = 0.004$). Also illustrated in the figure, the sum of positive ratings ('yes' to positive descriptions, plus 'no' to negative descriptions) was fewer for *Past Self* than any other individual ($F_{(3,69)} = 5.67$, $p = 0.002$) while no reliable difference was found among the ratings of *Present Self*, *Mother*, and *Queen*. These results are consistent with the well-established literature that self-esteem generally increases from adolescence to middle adulthood (e.g., Orth & Robins, 2014). However, a possible influence of behavioural performance was that motoric preparation and button reactions might contaminate neural decoding. If such performance-related factors indeed impacted on neural representations, it would be most manifest in the motor cortex and potentially spill over to other reas. To ascertain whether behavioural performance affected decoding results, we conducted control analyses using the patterns of primary motor cortex (Brodmann area 4). Standardised anatomical mask of the bilateral motor cortex was derived from the Wake Forest PickAtlas toolbox (Maldjian, Laurienti, Kraft, & Burdette, 2003) and warped into each participant's native psace using SPM's reverse-normalisation parameters. Using the nerual patterns of motor area, in three analyses we trained support-vector machines to differentiate *Present Self vs. Past Self*, *Mother vs. Queen*, and the four-way classification of different persons. As illustrated in the figure below, results showed that decoding accuracy did *not* differ from chance in any analysis (all $p > 0.24$). Crucially, the classifer could reliably differentiate *Mother* from *Queen* (see Figure 1 of the main article), despite the fact that the two conditions' reaction times and ratings were equated, effectively refuting these factors as alternative explnations. Together, these suggest that the decoding results could not simply be driven by performance-related variables.

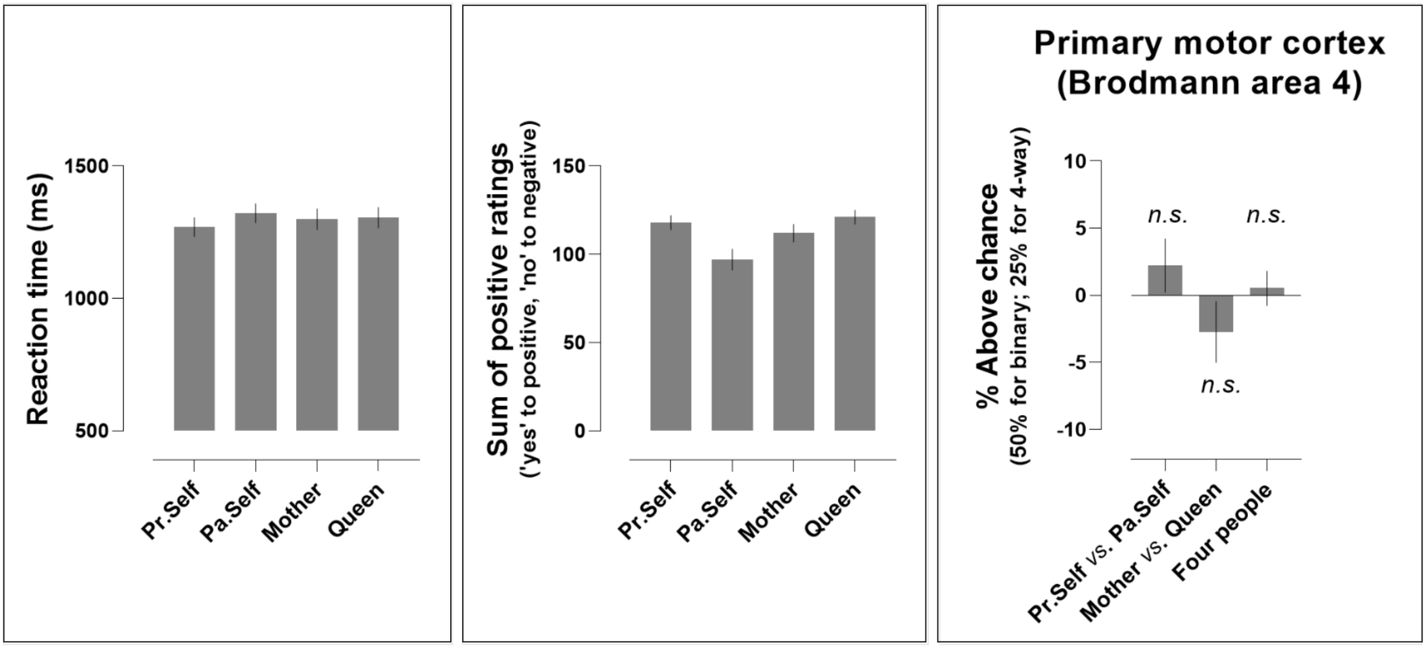

**Supplemental Results 2:** Decoding at the ROIs derived from meta-analyses

To ascertain that our findings do not depend the specific localiser paradigm of Dodell-Feder *et al.* (2011), we re-localised ROIs using the coordinates from three meta-analyses and performed multivoxel decoding at these literature-defined locations. For the semantic ROIs, we localised (1) the left and right anterior temporal lobes/ATLs based on Rice *et al.* (2015; see its Table 2; MNI: ±48, 8, -32) at the bilateral middle temporal gyrus and (2) the left inferior frontal gyrus/IFG based on Jackson (2020; see its Table 1) at the *pars opercularis* sub-region of the left IFG (MNI: -48, 20, 22). These two meta-analyses surveyed the fMRI literature on semantic processing and identified the bilateral ATLs and the left IFG as reliable neural regions for representing and retrieving semantic meaning, respectively. For the default-mode ROIs, we localised the dmPFC (MNI: -8, 56, 30), vmPFC (MNI: 0, 52, -12), PCC (MNI: 2, -56, 30), left IPL (MNI: -48, -56, 24), and right IPL (MNI: 56, -50, 18) using the meta-analysis of Bzdok *et al.* (2012; see its Table 1). These authors surveyed the literature of empathy and mentalising, and focused on the default-mode system. We identified corresponding points to these literature-defined coordinates in each person's native space and performed multivoxel-decoding to replicate our key results: (1) whether we could decode Present Self *vs.* Past Self, Mother *vs.* Queen, and cross-classify Self *vs.* Other across social distance and (2) whether decoding would be reliably better in the default-mode regions than semantic regions. Results below showed that the decoding based on meta-analyses are highly consistent with those based on functional localiser. Specifically, we found (1) while the ROIs were localised using a different approach, statistically robust above-chance decoding could still be detected in both semantic and default-mode ROIs and (2) decoding accuracy was consistently higher in the default-mode regions than in the semantic regions. Taken together, these additional data replicate our key results, support our conclusion, and demonstrate the robustness of our findings that they are not reliant on a specific functional localiser paradigm.

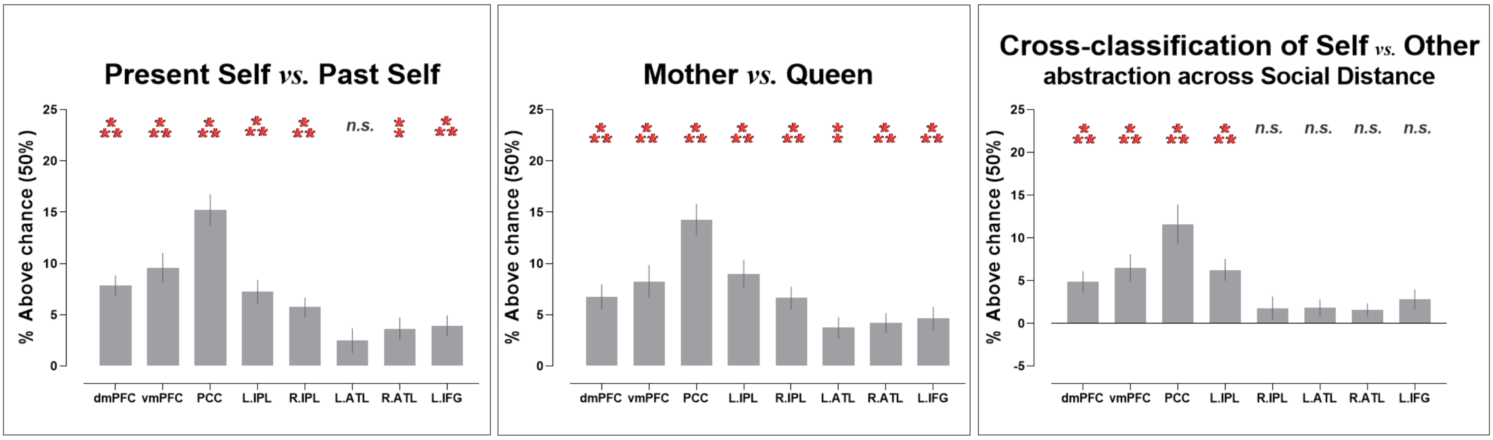

**Supplemental Results 3:** Comparison with NeuroSynth-defined semantic activation

The Dodell-Feder *et al.* (2011) localiser paradigm entails contrasting the Social against the Non-social condition to identify ROIs. Although this contrast has been able to identify clusters both in the default-mode network and in the semantic network (seen both in our data and the original study), the design of this contrast is inherently ‘social’ rather than ‘semantic’. To ensure that the three semantic ROIs (the left/right ATLs and the left IFG) identified with this approach are reliably engaged by semantic processing, we used NeuroSynth to identify the voxels that are robustly activated by semantic tasks (based on 40,030 activations from 1,031 neuroimaging studies), and compared our localiser results with the NeuroSynth results. As shown in the figure below, we found that the localiser contrast was able to identify all of the three semantic ROIs, and their locations were situated within the semantically-triggered clusters of NeuroSynth (localiser clusters: blue, NeuroSynth clusters: red, overlap: magenta), indicating that our localisation of semantic ROIs concur with the neurolinguistics literature.

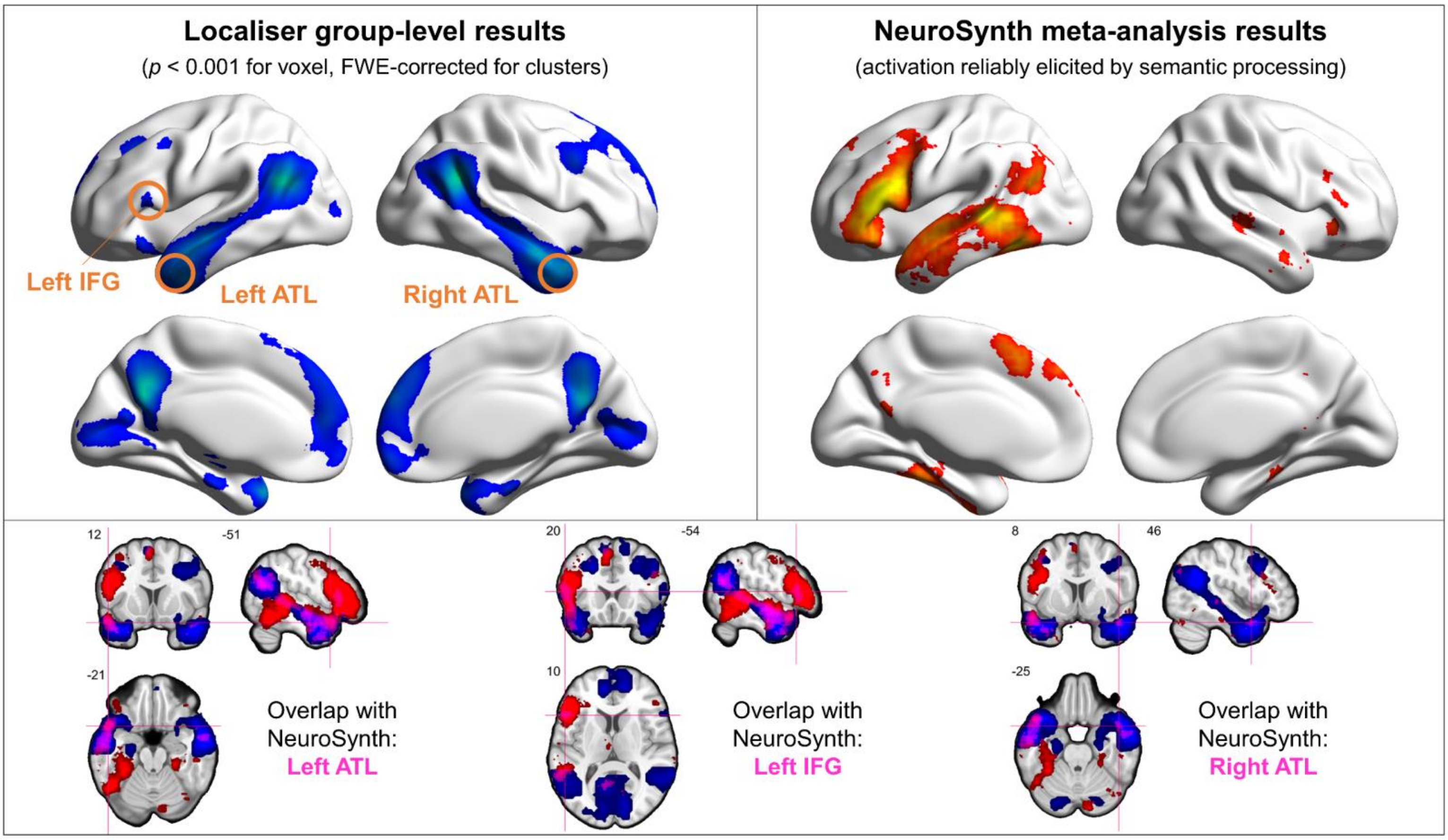

**Supplemental Results 4:** Comparison with NeuroSynth-defined self-related activation

While the Dodell-Feder *et al.* (2011) localiser paradigm was designed to probe ‘theory of mind’ (understanding another individual’s mental state) and the neural basis of mentalising, it also activated various regions involved in self-referential processing. We examined the pattern of activation in the original study (Dodell-Feder *et al.*, 2011), compared it with previous studies of self-referential tasks, and found much overlap of activation in the two threads of research. This overlap was also observed in our own empirical results – as shown in the figure below, when comparing our localiser data with the NeuroSynth results (based on 4,728 activations from 166 studies about self-referential processing), substantial overlap was found in various regions of the default-mode system (the dmPFC, vmPFC, PCC, as well as the bilateral IPL; localiser clusters: blue, NeuroSynth clusters: green, overlap: cyan). Taken together, the agreement between our ROIs and meta-analyses suggests that the Dodell-Feder paradigm provides an effective tool to identify broad swaths of areas in the default/semantic networks that are generally tuned to social cognition. Although there might be subtle variation between areas in terms of their tuning (e.g., the vmPFC prefers ‘self’ to ‘other’ but still reliably responds to ‘other’), the localiser has been highly effective to activate the whole system that has a general proclivity to social cognition (including both ‘self’ and ‘other’).

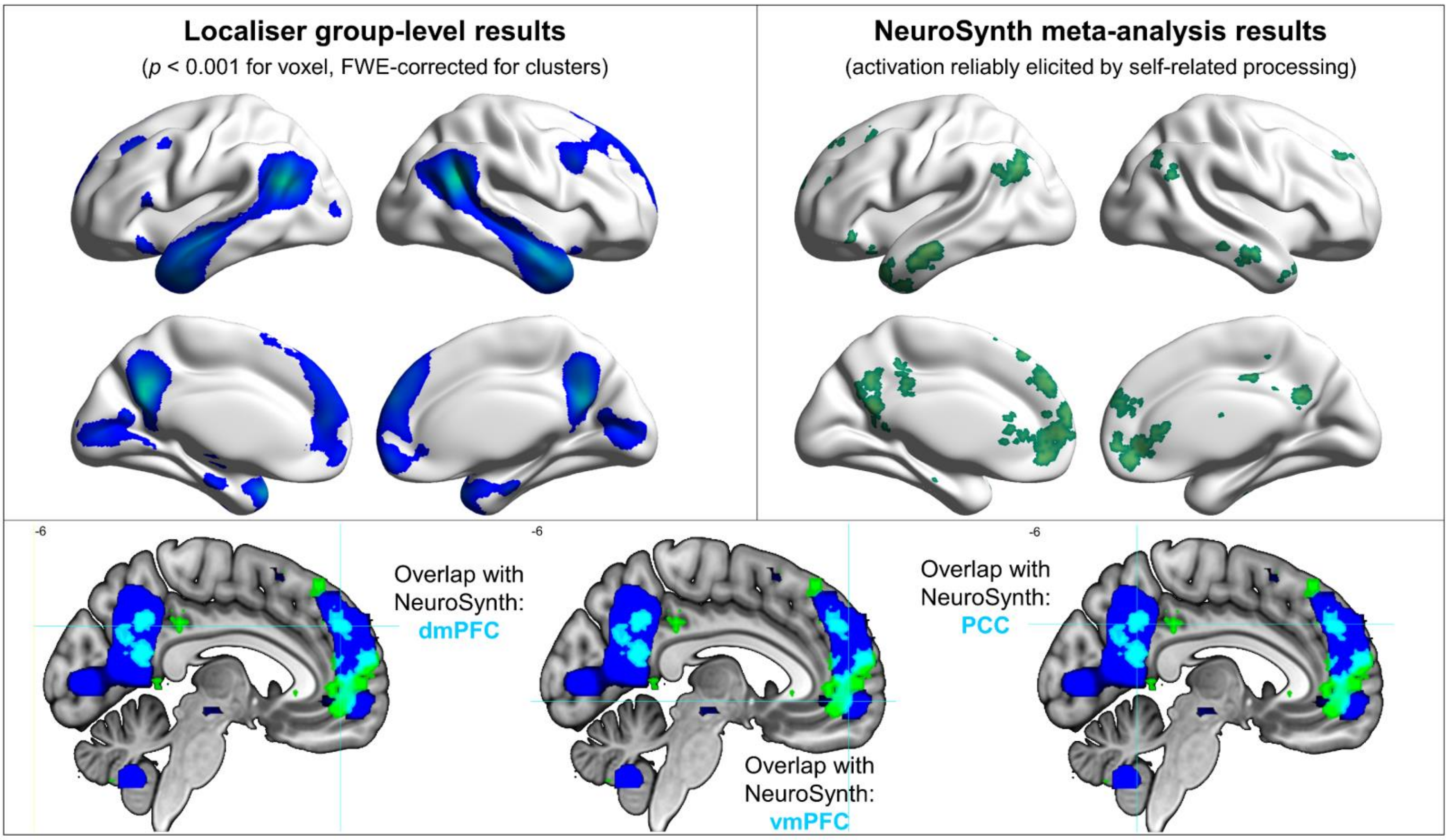

**Supplemental Results 5:** Whole-brain search for univariate effects

In addition to the multivariate analyses, we examined the univariate effects for three sets of contrast between conditions: Self (present & past) *vs.* Other (mother & Queen), Present Self *vs.* Past Self, Mother *vs.* Queen. Statistical threshold was $p < 0.001$ at the voxel-wise level, and FWE-correction ($p < 0.05$) for multiple comparisons at the cluster-level. Statistically reliable supra-threshold clusters were detected only in the following two comparisons: First, in the contrast of Present Self *vs.* Past Self, the rostral-medial prefrontal cortex showed greater activation for Present Self than Past Self, as indicated by the cluster in Figure (A). Second, in the comparison of Mother *vs.* Queen, three clusters were more active for Queen than Mother. As illustrated by Figure (B), these clusters are part of the default-mode network: the left/right inferior parietal lobules, and the right superior frontal gyrus.

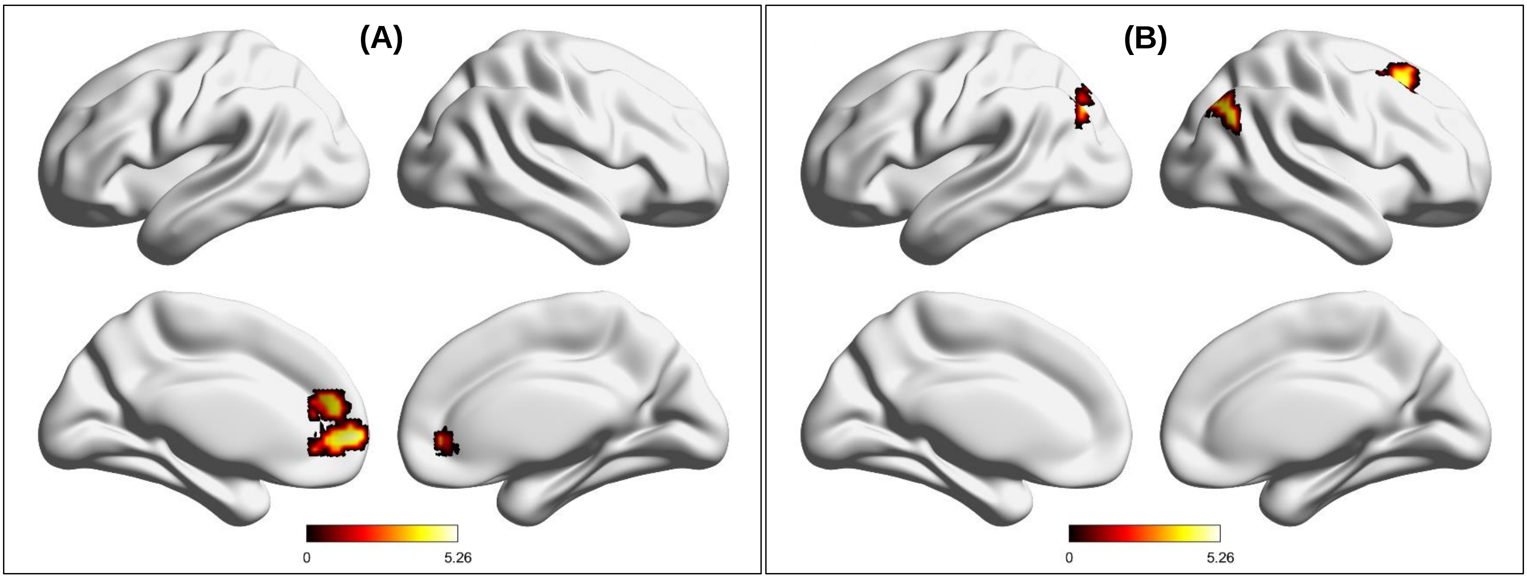

**<u>Present Self > Past Self</u>**

| Location | Cluster size | Peak (*t*-value) | MNI Coordinate (*x*, *y*, *z*) | | |
|---|---|---|---|---|---|
| Rostral-medial prefrontal cortex | 258 | 5.26 | 0 | 47 | 1 |

**<u>Queen > Mother</u>**

| Location | Cluster size | Peak (*t*-value) | MNI Coordinate (*x*, *y*, *z*) | | |
|---|---|---|---|---|---|
| Right inferior parietal lobule | 209 | 5.39 | 51 | -67 | 31 |
| Left inferior parietal lobule | 203 | 5.32 | -36 | -79 | 43 |
| Right superior frontal gyrus | 165 | 4.88 | 27 | 23 | 46 |

**<u>Past Self > Present Self</u>, <u>Mother > Queen</u>, <u>Self > Other</u>, <u>Other > Self</u>**
No supra-threshold clusters for these contrasts

**Supplemental Results 6:** Confusion matrix

In this supplementary analysis, we report confusion matrices for the 4-way classification of four target individuals (*Present Self*, *Past Self*, *Mother*, and *Queen*; Figure 1A of main article), computed separately for each region of interest. The vertical axis of a matrix indicates the ground truth (actual labels), whereas the horizontal axis indicates the label predicted by the SVM classifier. The value in each cell indicates the percentage of prediction for a given pair (e.g., % of predicting the 'Mother' label when the actual label was indeed 'Mother'). The colour-coding provides a visual layout of the classifier's performance (darker/lighter blue – lower/higher percentage). We observed a highly consistent pattern across all eight regions of interest. Illustrated below are the example matrices of the posterior cingulate cortex and ventromedial prefrontal cortex. It is noteworthy that the analysis shown here is a standard *cross-validation* decoding, rather than the *cross-classification* of four individuals across emotive valence (Figure 3B of main article). However, the two analyses yielded entirely coherent results – when decoding a person's identity, the algorithm was most confused with a socially proximal concept (e.g., mistaking *Present Self* as *Past Self* – more probable error) and least confused with a socially distant concept (e.g., mistaking *Present Self* as *Queen* – least probable error).

Posterior cingulate cortex

| | PrSelf | PaSelf | Mother | Queen |
|---|---|---|---|---|
| PrSelf | 46 | 24 | 17 | 12 |
| PaSelf | 24 | 45 | 13 | 17 |
| Mother | 23 | 15 | 41 | 20 |
| Queen | 17 | 18 | 20 | 45 |

Ventromedial PFC

| | PrSelf | PaSelf | Mother | Queen |
|---|---|---|---|---|
| PrSelf | 43 | 22 | 19 | 17 |
| PaSelf | 29 | 39 | 14 | 18 |
| Mother | 27 | 20 | 32 | 21 |
| Queen | 22 | 24 | 23 | 31 |
